# A versatile pattern-forming cortical circuit based on Rho, F-actin, Ect2, and RGA-3/4

**DOI:** 10.1101/2022.03.08.483353

**Authors:** Ani Michaud, Marcin Leda, Zachary T. Swider, Songeun Kim, Jiaye He, Jennifer Landino, Jenna R. Valley, Jan Huisken, Andrew B. Goryachev, George von Dassow, William Bement

## Abstract

Many cells can generate complementary traveling waves of actin filaments (F-actin) and cytoskeletal regulators. This phenomenon, termed cortical excitability, results from coupled positive and negative feedback loops of cytoskeletal regulators. The nature of these feedback loops, however, remains poorly understood. We assessed the role of the Rho GAP RGA-3/4 in the cortical excitability that accompanies cytokinesis in both frog and starfish. RGA-3/4 localizes to the cytokinetic apparatus, “chases” Rho waves in an F-actin-dependent manner and, when co-expressed with the Rho GEF Ect2, is sufficient to convert the normally quiescent, immature *Xenopus* oocyte cortex into a dramatically excited state. Experiments and modeling show that changing the ratio of RGA-3/4 to Ect2 produces a range of cortical behaviors from pulses to complex waves of Rho activity. We conclude that RGA-3/4, Ect2, Rho and F-actin form the core of a circuit that drives a diverse range of cortical behaviors, and demonstrate that the immature oocyte is a powerful model for characterizing these dynamics.

**Summary:** Michaud et al. identify Ect2 and RGA-3/4 as core components of the cortical excitability circuit associated with cytokinesis. Additionally, they demonstrate that the immature *Xenopus* oocyte is a powerful model for characterizing excitable dynamics.

## Introduction

The cell cortex interprets and responds to a wide variety of intra- and extracellular cues by forming dynamic patterns of cytoskeletal proteins that accomplish local changes in cell shape. For example, during cytokinesis the cortical response to signals arising from the mitotic spindle is to assemble the cytokinetic apparatus, an equatorial array of F-actin and myosin-2 that drives an ingressing constriction. During chemotaxis, the cortical response to chemoattractant-receptor binding is to reorganize the cortical cytoskeleton locally such that F-actin assembly and disassembly result in cortical protrusions and retractions that accomplish cell movement up the chemoattractant gradient.

The Rho GTPases – Rho, Rac and Cdc42 – are key regulators of cytoskeletal assembly that mediate many of the cell’s cortical behaviors. These small switch-like proteins cycle between an active, GTP-bound state, and an inactive GDP-bound state (Etienne-Manneville and Hall, 2002). Cycling by the intrinsic GTPase activity is so slow that dynamic behaviors by the Rho family depend on regulators that promote nucleotide exchange (Guanine nucleotide exchange factors; GEFs) and hydrolysis (GTPase activating proteins; GAPs; Goryachev and Pokhilko, 2006). In contrast to a classical view of these signals as whole-cell “state switches”, live-cell visualization of Rho GTPase activity shows that active GTPases are commonly deployed in distinct, local, and dynamic patterns – patches, stripes, rings, and waves (Bement et al., 2006). These patterns contribute significantly to cell behavior by recruiting effector proteins that modulate F-actin and myosin-2. Thus, the Rho GTPases and their effectors sculpt the physical structure, shape, and motion of the cortex on the timescale of seconds to minutes.

One of the most fascinating examples of cortical pattern formation is cortical excitability, a self-organized behavior characterized by traveling waves of F-actin, often under the control of complementary waves of GTPase activity (Bement et al., 2015). Cortical excitability has been observed in many different cell types and cellular processes (Barnhart et al., 2017; Gerhardt et al., 2014; Graessl et al., 2017; Michaux et al., 2018; Pal et al., 2019; Weiner et al., 2007; Xiao et al., 2017), and though the underlying control networks differ, the key features of excitable signaling are thought to be similar: fast, positive feedback at the wave front is responsible for wave propagation, while delayed, negative feedback at the trailing edge of the wave transitions the system to a refractory state.

Although there is long-standing consensus on the basic conceptual scheme of cortical excitability, characterization of the actual circuit participants and their relationships during real cell behaviors of interest is challenging and remains incomplete. Cortical excitability during cytokinesis offers a case in point: in starfish and frog embryos, waves of Rho activity and F-actin polymerization traverse the cortex and are focused at the equator by the anaphase spindle (Bement et al., 2015). Rho activation is thought to be amplified by a positive feedback loop involving Ect2 (Chen et al., 2020), while delayed negative feedback is somehow linked to F-actin (Bement et al., 2015). Completing this fragmentary scheme is complicated by the fact that anaphase spindle configuration and Ect2 activity are critically dependent on cell cycle progression (Glotzer, 2009; Green et al., 2012; Tatsumoto et al., 1999) and limited by the transience of cytokinesis-phase (C-phase; Canman et al., 2000). Further complexity stems from the fact that other cytokinetic participants modulate Ect2 distribution and activity (Frenette et al., 2012; Kim et al., 2014; Tatsumoto et al., 1999; Zhang and Glotzer, 2015), and can influence Rho activity independently of Ect2 (Budnar et al., 2019; Miller and Bement, 2009; Su et al., 2003). These complications make it desirable to isolate the circuit that mediates excitability and evoke its activity at steady state.

Here we investigate the role of the Rho GAP RGA-3/4 (also known as ArhGAP11a/MP-GAP) in cortical excitability and developed a near-steady-state reconstitution of cortical excitability in immature oocytes. We find that RGA-3/4—previously shown to negatively regulate Rho during cytokinesis (Bell et al., 2020; Schmutz et al., 2007; Schonegg et al., 2007; Zanin et al., 2013) and pulsed contractions in *C. elegans* (Michaux et al., 2018)— localizes to the cytokinetic apparatus in both starfish and frog embryos and participates in excitable dynamics by negatively regulating Rho. We further show that co-expression of RGA-3/4 and Ect2 is sufficient to induce high-level cortical excitability in immature frog oocytes, which are normally cortically quiescent. This oocyte system can be tuned to display a remarkably rich range of dynamic cortical behaviors by modulating the Ect2/RGA-3/4 ratio, and therefore represents a simple yet faithful system for investigating cortical excitability during cytokinesis. These behaviors are captured by a theoretical model based on positive feedback involving Rho and Ect2, and delayed negative feedback involving Rho, RGA-3/4 and F-actin. Our results demonstrate that Ect2 and RGA-3/4, along with F-actin and Rho, form the core of a conserved cortical excitability circuit involved in cytokinesis.

## Results

### Starfish RGA-3/4 recruits to and modulates cortical waves

Because the dynamic localization of RGA-3/4 has only been described for pulsed contractions in *C. elegans* embryos (Bell et al., 2020; Michaux et al., 2018), we first characterized RGA-3/4 during meiosis and mitosis in starfish (*P. miniata*) using mNeon fused with either wild-type RGA-3/4 (mNeon-RGA-3/4^WT^) or GAP-dead RGA-3/4 (mNeon-RGA-3/4^R96E^) from *P. miniata*. Both wild-type and GAP-dead derivatives localized to the cytokinetic furrow during meiotic and embryonic mitotic divisions (Fig. 1A, B and Fig. S1B, D). However, when expressed at levels needed for clear visualization, wild-type RGA-3/4 reduced Rho activity (see below), making it necessary to use GAP-dead RGA-3/4 in situations where normal Rho activity was required (i.e. cytokinesis).

**Figure 1.**
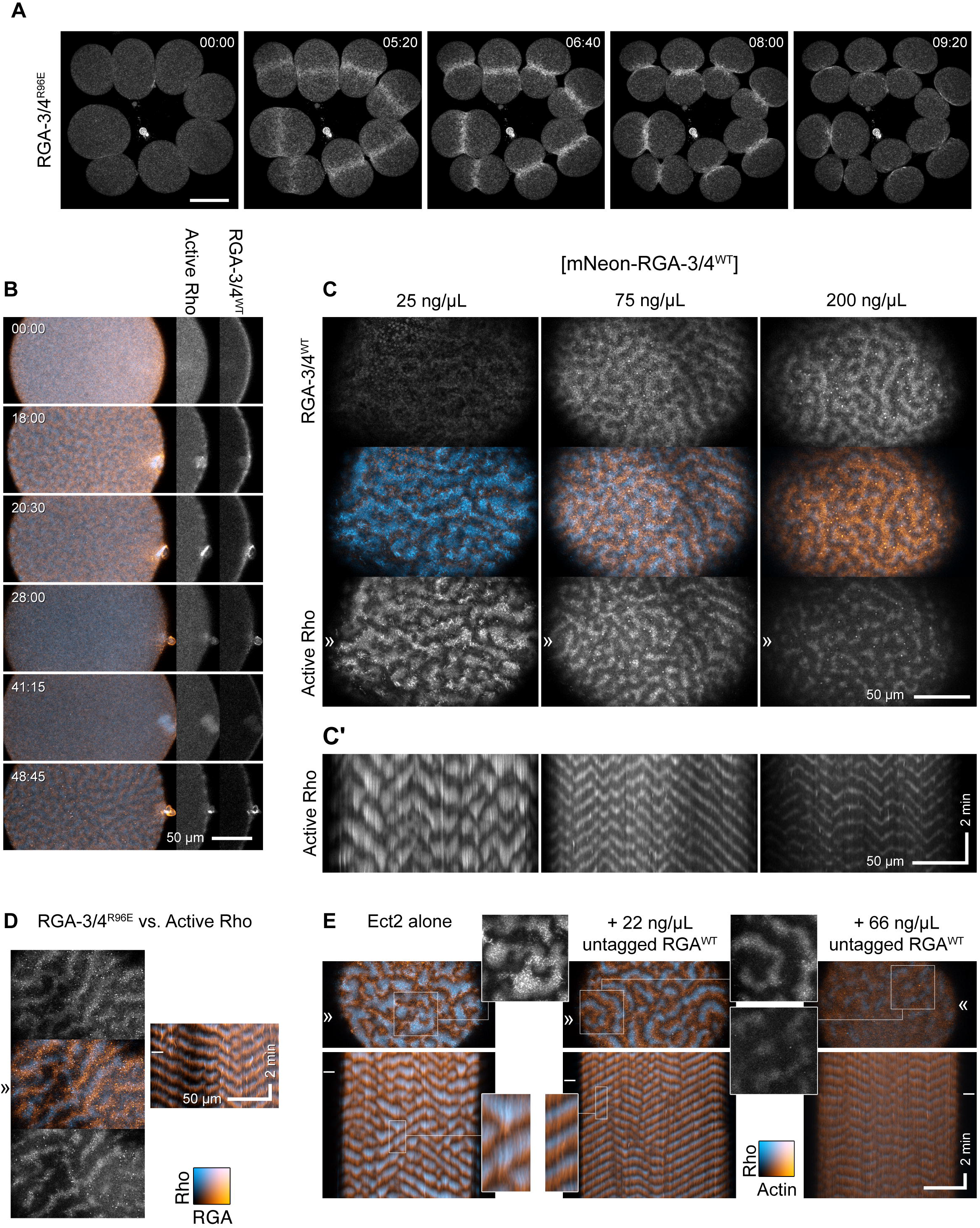
RGA-3/4 localizes to the cortex, cortical waves, the cytokinetic apparatus, and suppresses Rho activity in starfish eggs and embryos. (**A**) Time course of mitosis in starfish blastomeres (8 of 16 cells) expressing mNeon-RGA-3/4^R96E^. mNeon-RGA-3/4^R96E^ first localizes to the cortex, then the equatorial cortex, then the cytokinetic apparatus. Time in min:sec. (**B**) Time course of an Ect2-loaded starfish oocyte undergoing first and second meiosis; animal pole is to the right, time in min:sec, active Rho labeled with mCherry-rGBD (cyan) and mNeon-RGA-3/4^WT^ (orange). Waves develop coincident with polar body emission and subside between meiosis I and II. Single-channel insets of the animal pole use the deepest slice to show cortical recruitment: RGA-3/4 is cortical as the cell approaches meiosis-I (MI) metaphase, while Rho is not (00:00); RGA-3/4 appears brightly in the polar body furrow (frame 3); RGA-3/4 departs the cortex between MI and MII (frame 4) but returns in metaphase (frame 5) before waves develop. (**C**) Waves of wild-type RGA-3/4 (mNeon-RGA-3/4^WT^; orange) recruitment closely follow Rho activity waves (mCherry-rGBD; cyan), and increasing dose of mNeon-RGA-3/4^WT^ progressively suppresses Ect2-induced excitability (Video 1); kymographs in C’ are taken from the band denoted by “»”. All panels show post-MII oocytes at quasi-steady state. All oocytes expressing 100 ng/μl Ect2 to induce rampant, chaotic waves. Those simultaneously loaded with 25 ng/μl mNeon-RGA-3/4 are little different from controls (not shown); 75 ng/μl or 200 ng/μl RGA-3/4 reduces wave amplitude and peak width, while extending wave propagation into long runs. (**D**) GAP-dead RGA-3/4 (mNeon-RGA-3/4^R96E^; orange) co-expressed with Ect2 has no effect on excitability, but recruits in the same phase as wild-type RGA-3/4 (see also, Video 2); active Rho (mCherry-rGBD; cyan). Kymograph position corresponds to “»”, stills come from the time indicated by “–”. Color swatch applies to panels B-D. (**E**) Untagged wild-type RGA-3/4 (RGA-3/4^WT^) co-expressed with high level (100 ng/μl) Ect2, labeled with GFP-rGBD (active Rho; cyan) and mCherry-UtrCH (F-actin; orange). Corresponds to Video 3. In Ect2 alone samples (left panel), waves are irregular, close-packed high-amplitude bursts that form broken fronts that swell and collapse; addition of modest (22 ng/μL) RGA-3/4 mRNA converts them to steadily-rolling regular waves (middle panel); higher RGA-3/4 dose damps waves further (right panel), and higher still suppresses them completely (not shown). Insets from stills are 2x blowups of Rho alone. Insets from kymographs are 3x blowups.

RGA-3/4 localized to the germinal vesicle (nucleus) of immature starfish oocytes (Fig. S1A). Following treatment with 1-methyladenine to induce meiotic maturation, RGA-3/4 was released into the cytoplasm at germinal vesicle breakdown and thereafter associated with the cortex prior to meiosis I and immediately before meiosis II (Fig. 1B). During polar body emission (meiotic cytokinesis), RGA-3/4 accumulated prominently in the cleavage furrow (Fig. 1B, Fig. S1B). Notably, during mitosis, RGA-3/4 localized transiently to the entire cortex before becoming restricted equatorially (Fig. S1C, D). To determine the localization of RGA-3/4 during cortical excitability, we targeted the post-meiotic period in starfish oocytes, during which a sufficient dose of extra Ect2 elicits sustained propagation of steady waves over 1-2 hours (see methods and Bement et al., 2015). Strikingly, in starfish oocytes overexpressing Ect2 and a probe for active Rho (rGBD; Benink and Bement, 2005), wild-type and GAP-dead mNeon-RGA-3/4 associated with the cortex in waves, with a peak that followed Rho activity peaks by 15.7 ± 1.9 seconds (n=7 cells) (Fig. 1C, D, Fig. S1E, Video 1).

The dynamic localization of RGA-3/4—in meiotic and mitotic furrows and behind waves of Rho activity—is consistent with a role in negative feedback. Such a role predicts that increasing RGA-3/4 expression should attenuate Rho activity. To test this, we amplified cortical excitability by loading starfish oocytes with a moderately high dose of Ect2 (see Methods and Bement et al., 2015) and increasing doses of tagged, wild-type RGA-3/4 (mNeon-RGA-3/4^WT^ Fig. 1C). The selected dose of Ect2 was sufficient to elicit high-amplitude, chaotic cortical Rho waves after oocytes had completed meiosis. At low concentrations, wild-type mNeon-RGA-3/4 had no detectable effect on Rho wave amplitude or wave form, even though faint waves of RGA-3/4 recruitment were apparent above the background autofluorescence of yolk (Fig. 1C, 25 ng/μL). At higher concentrations, wild-type mNeon-RGA-3/4 had a profound effect on both wave amplitude and form: chaotic, heaping wave bursts were converted to low-amplitude, steadily-propagating swells (Fig. 1C, 75 ng/μL; Video 2). This transformation is most evident in kymographs (Fig. 1C’), wherein the large-amplitude bursts in the lowest dose are replaced by long, steady traces in the middle dose. At the highest dose, tagged, wild-type RGA-3/4 further suppressed Rho wave amplitude (Fig. 1C, 200 ng/μL and C’).

We also evaluated untagged wild-type RGA-3/4 and found that it behaves in every way similarly to wild-type mNeon-RGA-3/4, but is noticeably more potent: when co-expressed with a high dose of Ect2 along with probes for active Rho (GFP-rGBD) and F-actin (mCherry-UtrCH; Burkel et al., 2007), increasing wild-type RGA-3/4 levels regularized propagating wavefronts and reduced amplitude without altering the relationship between Rho activity and actin assembly that is the basis of the cortical wave cycle (Fig. 1E, Video 3). Quantification of the impact of RGA-3/4 expression revealed that it reduced wave period (Fig. S1F), temporal width (Fig. S1G), and dramatically reduced wave amplitude (Fig. S1H). In contrast, the GAP-dead point mutant (mNeon-RGA-3/4^R96E^) had no impact on Rho wave dynamics or wave duration at any concentration tested (Fig. 1D and not shown). Together, these results show that RGA-3/4 negatively regulates excitable Rho activity during starfish cytokinesis.

### RGA-3/4 behaves like an actin-recruited Rho inhibitor

RGA-3/4 localization has been linked to F-actin polymerization (Michaux et al., 2018), and previous work has shown that F-actin is involved in negatively regulating Rho activity during cortical excitability (Bement et al., 2015). We therefore next sought to investigate the relationship between RGA-3/4 and F-actin. In Ect2-expressing starfish oocytes, mNeon-RGA-3/4^WT^ colocalizes with F-actin, occupying almost exactly the same phase of the wave cycle (Fig. 2A). However, cross-correlational analysis revealed a slight lead of RGA-3/4 recruitment with respect to peak F-actin signal (4.2 ± 2.4 seconds; n=4 cells) (Fig. 2B).

**Figure 2.**
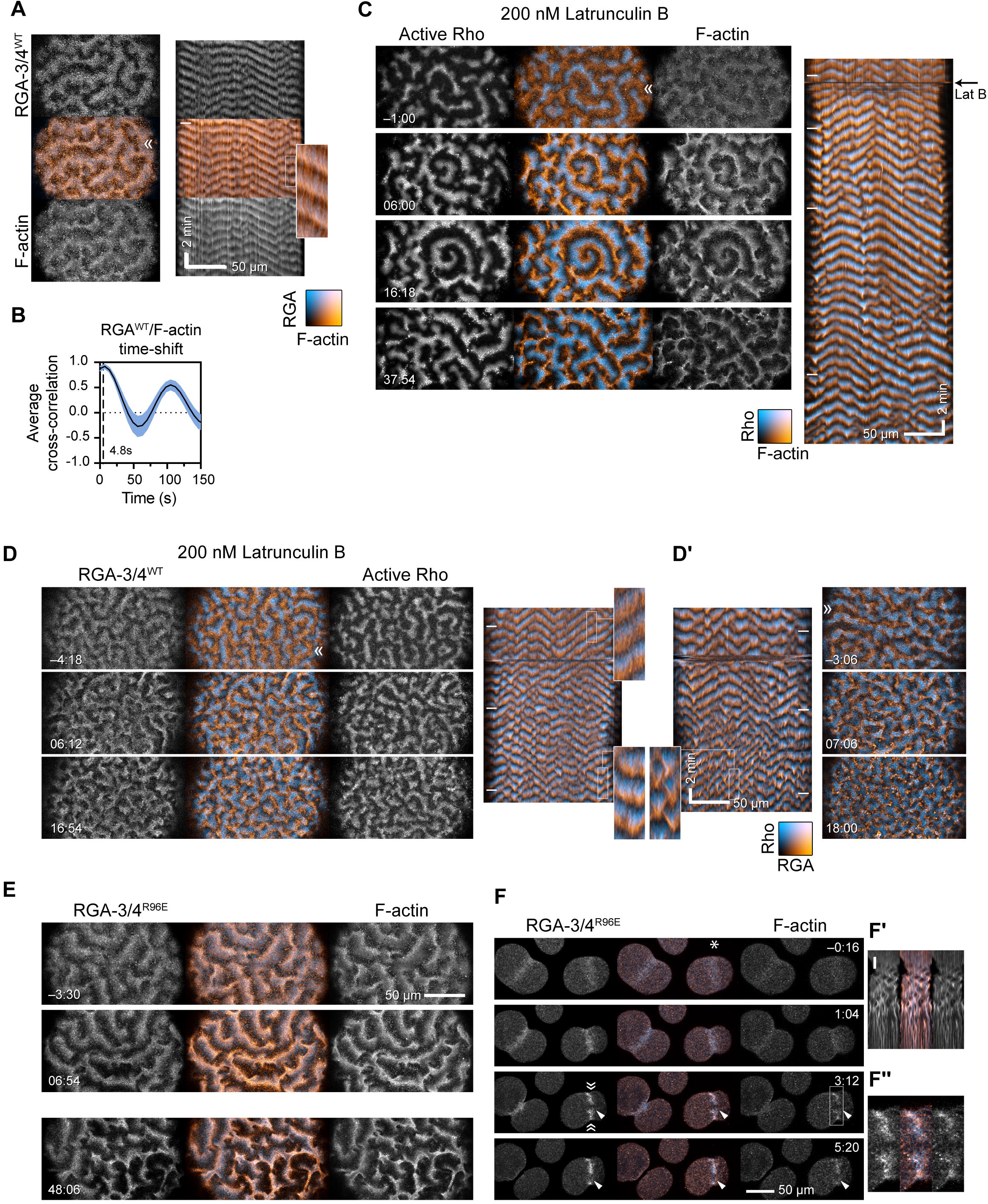
Starfish RGA-3/4 behaves like an actin-dependent Rho inhibitor. (**A**) Stills and corresponding kymograph from a post-MII oocyte co-expressing Ect2 (100 ng/μl) and mNeon-RGA-3/4^WT^ (cyan; 75 ng/μl) as well as mCherry-UtrCH (orange) to detect F-actin. RGA-3/4 and F-actin overlap almost perfectly. RGA-3/4 image and kymograph is from maximum intensity projection of 3 z-slices per timepoint, minus 0.9 times the minimum value over a 20 timepoint sliding window to reduce background autofluorescence. Kymograph position corresponds to “»“. (**B**) Cross-correlational analysis of a representative starfish cell expressing mNeon-RGA-3/4^WT^ and mCherry-UtrCH showing a 4.8 s delay between peak RGA recruitment peak actin signal. **(C)** Rho activity (GFP-rGBD; cyan) versus F-actin (mCherry-UtrCH; orange) in a post-MII Ect2-loaded oocyte flooded with 200 nM Latrunculin B at time 0 (corresponds to Video 4). Before treatment this cell experienced steady high-amplitude rolling waves; within minutes of treatment, wave amplitude is noticeably enhanced. Black arrow to right of kymograph indicates Latrunculin B addition at time 0. (**D**) Same experiment as in (C) but with mNeon-RGA-3/4^WT^ (orange) instead of UtrCH (corresponds to Video 5). Dose of Ect2 and RGA-3/4 titrated to induce steady, rolling waves before treatment. After treatment, Rho amplitude is noticeably enhanced and waves are more closely packed. RGA-3/4 continues to occupy a phase immediately following Rho. (**D’**) Another oocyte from the same batch, in which the pre-treatment behavior is somewhat higher on the excitability spectrum. Treatment likewise enhances wave amplitude and packs waves more tightly, and breaks wave fronts into irregular bursts. RGA-3/4 images background-subtracted as in (A). **(E)** GAP-dead RGA-3/4 continues to track F-actin throughout latrunculin treatment. Similar treatment to C,D but with mNeon-RGA-3/4^R96E^ (left; cyan) and mCherry-UtrCH (right; orange). Times are min:sec relative to flooding with 200 nM Latrunculin B. RGA-3/4 images are not background-subtracted. Top two panels are from one oocyte, the bottom one from another oocyte in the same treatment batch. Color swatch in A applies to B. **(F)** Cleaving embryonic cells co-expressing mNeon-RGA-3/4^R96E^ and mCherry-UtrCH, the latter underlabeled to avoid interfering with actin-dependent events; a micropipette filled with 0.5% LM agarose + 4 μM Latrunculin B is parked next to the cell on the right and moved into position (white asterisk) at time 0. Furrow stalls within minutes; F-actin band breaks into pulsed contractions (kymograph, inset F’, generated from position “» «” – compare to Fig. S1C); RGA-3/4 continues to nearly match F-actin (2.5× blowup, inset F”).

If RGA-3/4 recruitment is linked to actin polymerization, then delaying or reducing actin assembly should a) commensurately alter RGA-3/4 recruitment to the cortex while b) reversing the effect of RGA-3/4 on wave characteristics. Extreme reduction of F-actin is of no help here as this simply results in a burst of cortical Rho activity followed by cessation of wave propagation (Bement et al., 2015). We therefore titrated the concentration of the actin assembly inhibitor, latrunculin B, to find a concentration that altered F-actin distribution without eliminating excitability. We found that 200 nM latrunculin B reliably induced a noticeable narrowing of the F-actin waves, which was accompanied by a brightening and broadening of the Rho waves. This treatment also converted steady waves to more chaotic, broken bursts (Fig. 2C, and Video 4).

To assess the effects of latrunculin treatment on RGA-3/4 recruitment, we loaded oocytes with a combined dose of Ect2 and wild-type mNeon-RGA-3/4 titrated both to visualize RGA-3/4 recruitment and to produce steady, rolling waves, and again treated with 200 nM latrunculin B. As above, the latrunculin treatment resulted in a brightening and broadening of the Rho waves. Consistent with a link between F-actin and RGA-3/4, latrunculin treatment resulted in a narrowing of the RGA-3/4 waves (Fig. 2D, D’). To ensure that the observed behavior of mNeon-RGA-3/4^WT^ following latrunculin treatment did not reflect the combined impact of actin manipulation and RGA-3/4-mediated alterations in excitable behavior, we repeated the latrunculin treatment using GAP-dead mNeon-RGA-3/4, which exhibited the same behavior as wild-type mNeon-RGA-3/4 (Fig. 2E). Finally, to assess the relationship between F-actin and RGA-3/4 in the absence of Ect2 overexpression, we took advantage of the fact that the cytokinetic apparatus can be fractured by localized treatment with latrunculin using a micropipet (Bement et al., 2015). As shown in Fig. 2F, focal latrunculin treatment of cleaving embryonic cells stalled furrow ingression and induced cycles of contraction within the damaged cytokinetic apparatus; during these cycles RGA-3/4 and F-actin remained extensively colocalized.

### RGA-3/4 dynamics in frog embryos

The results outlined above indicate that starfish RGA-3/4 participates in actin-dependent negative feedback, first engaging newly-assembled F-actin elicited by active Rho, then terminating Rho autoactivation to return Rho activity to baseline levels. Activated eggs and embryonic cells of the frog *Xenopus laevis* exhibit qualitatively similar Rho-dependent excitability but do so even without exogenous Ect2 and during a broader fraction of the cell cycle (Bement et al., 2015). RNA encoding wild-type *Xenopus* RGA-3/4 fused to three tandem copies of GFP (RGA-3/4^WT^-3xGFP) caused cytokinesis defects at expression levels required for imaging early developmental timepoints (data not shown). However, we were able to visualize the protein at later stages (~stage 9-10) by injecting an extremely low dose of mRNA and letting expression ramp up slowly. Imaging at earlier developmental stages was achieved using a GAP-dead mutant (RGA-3/4^R80E^-3xGFP), which could be expressed at higher levels.

Consistent with our findings in starfish, both wild-type and GAP-dead RGA-3/4 localized to the nucleus during interphase and concentrated at the cortex following nuclear envelope breakdown (Fig. 3A, B, Fig. S2; Video 5). Both also concentrated at the equatorial cortex prior to and during cytokinesis, where they colocalized with F-actin (Fig. 3A, B, C; Video 6). The higher levels of expression permitted by the GAP-dead mutant revealed low amplitude waves outside the equator and higher amplitude waves at the equator during cytokinesis (Fig. 3B, B’, C, C’, Video 7). Comparison of RGA-3/4 to an Factin probe (UtrCH) showed that while both are found in furrow and non-furrow waves, the ratio of RGA-3/4 to F-actin is higher at the equator than outside the equator (Fig. 3C, C’).

**Figure 3.**
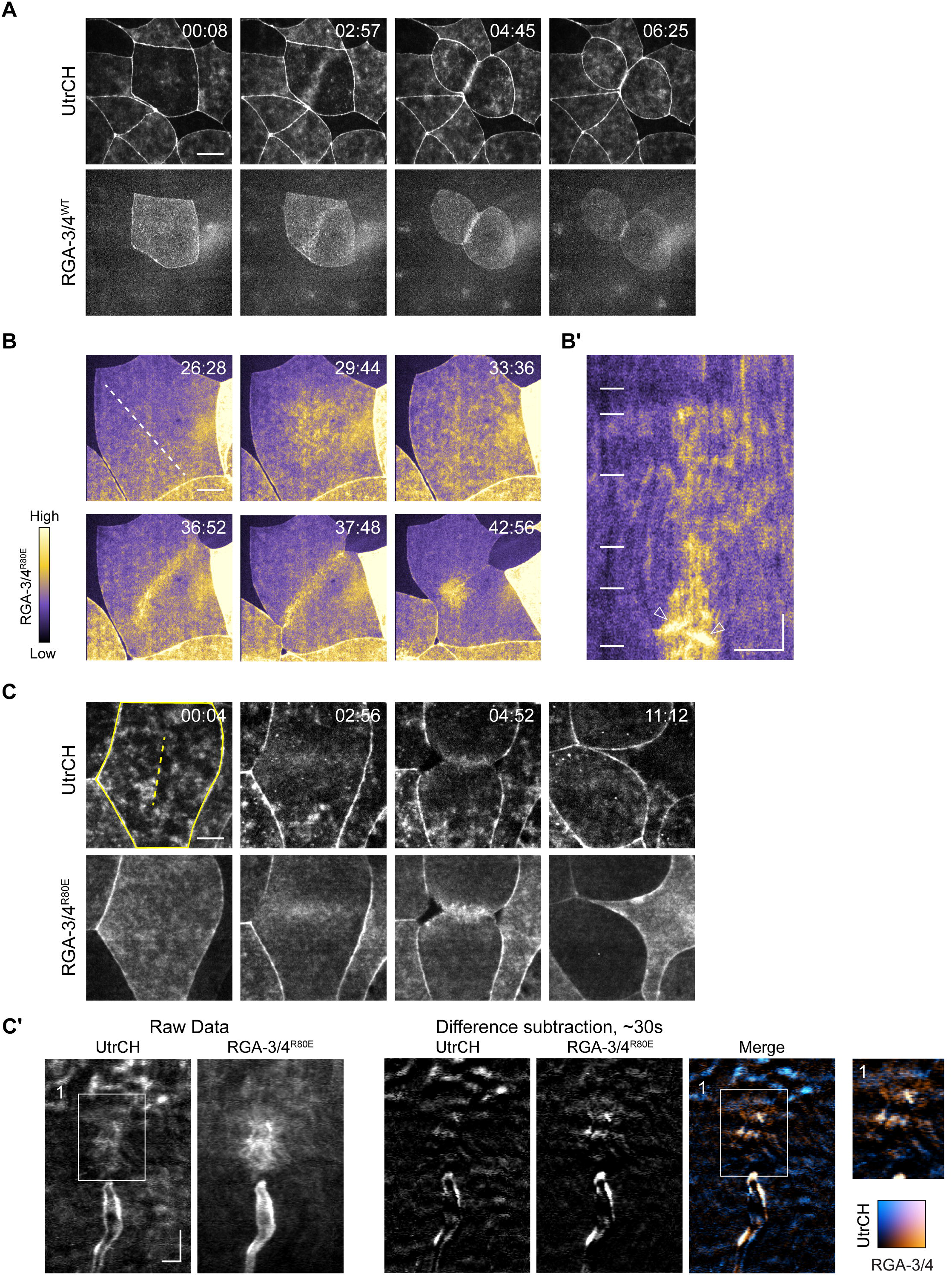
RGA-3/4 localizes to the equatorial cortex and contractile ring in *Xenopus* embryos. (**A**) Time course of F-actin and RGA-3/4 ^WT^-3xGFP in *Xenopus* embryo epithelial cell undergoing cytokinesis. Time in min:sec, F-actin (mCherryUtrCH, top) and tagged, wildtype RGA-3/4 (RGA-3/4 ^WT^-3xGFP, bottom). RGA-3/4 signal accumulates at equatorial cortex coincident with F-actin localization and persists throughout cytokinesis (see also, Video 6). Scale bar = 25 μm. (**B**) GAP-dead RGA-3/4 localizes to cortical waves, the equatorial cortex and contractile ring in *Xenopus* embryos. Time course of single cell in early frog embryo undergoing cytokinesis. Tagged, GAP-dead RGA-3/4 (RGA-3/4^R80E^-3xGFP) localizes to cortical waves (29:44, 33:36), equatorial cortex (36:52) and contractile ring (37:48 and 42:56); see also, Video 7. Scale bar = 25 μm. (**B’**) Kymograph generated from region indicated by dotted line in (B). Positions of still frames in (B) indicated on kymograph with white dashes. Waves in furrow region are labeled with white arrowheads. x scale bar = 25 μm, y scale bar = 2 min. (**C**) RGA-3/4 and actin co-localize in furrow waves. Time course of cell in dividing frog embryo expressing mCherry-UtrCH (actin; top) and RGA-3/4 ^R80E^-3XGFP (bottom). Scale bar = 25 μm. Kymographs (**C’**) correspond to dotted yellow line (panel 1). Kymographs generated from raw data (C’, left) and difference subtraction data (C’, right); Inset from box 1 is magnified 1.4x. x scale bar = 5 μm, y scale bar = 2 min.

### A combination of Ect2 and RGA-3/4 induces high amplitude waves in immature oocytes

Our previous results suggested that Rho autoactivation is mediated by Ect2, whereas the results above point to RGA-3/4 as a candidate for actin-dependent Rho autoinhibition. Modeling suggests that these two factors should suffice to sustain cortical excitability. To test this hypothesis, we sought to induce excitability in the cortex of a cell type that does not naturally display excitability, namely, the immature *Xenopus* oocyte (Bement et al., 2015). Consistent with previous results, immature oocytes expressing probes for active Rho (rGBD) displayed no evidence of cortical excitability (Fig. 4A). Overnight expression of non-importable Ect2 (Ect2^ΔNLS^; to prevent sequestration in the nucleus) had varied effects: in most cases, it elevated Rho activity without inducing waves, while in some cases it induced small patches of short, low-amplitude, pulses (Fig. 4B, Fig. S3A). In contrast, co-expression of Ect2^ΔNLS^ and wild-type RGA-3/4 induced waves throughout the cortex of a high proportion of the oocytes (up to 100% in some experiments) (Fig. 4C, Video 8). The waves induced by the combination of Ect2 ^ΔNLS^ and RGA-3/4 were high in amplitude and formed complex patterns with, in many cases, well-developed spiral waves that spanned a large expanse of the cortex (Fig. 4C). Overexpression of wild-type RGA-3/4 alone did not produce these patterns (Fig. 4D). Cells co-expressing Ect2^ΔNLS^ and wildtype RGA-3/4 also displayed waves with higher relative amplitudes (Fig. 4E) and longer, more connected wave-trains (Fig. S3B, C). The number of oocytes displaying cortical waves also increased from roughly 27% in cells expressing Ect2^ΔNLS^ alone – all of which manifested local low-amplitude waves only – to around 80% when both Ect2^ΔNLS^ and RGA-3/4 were present, typified by coherent, propagating fronts across large spatial domains (Fig. S3D).

**Figure 4.**
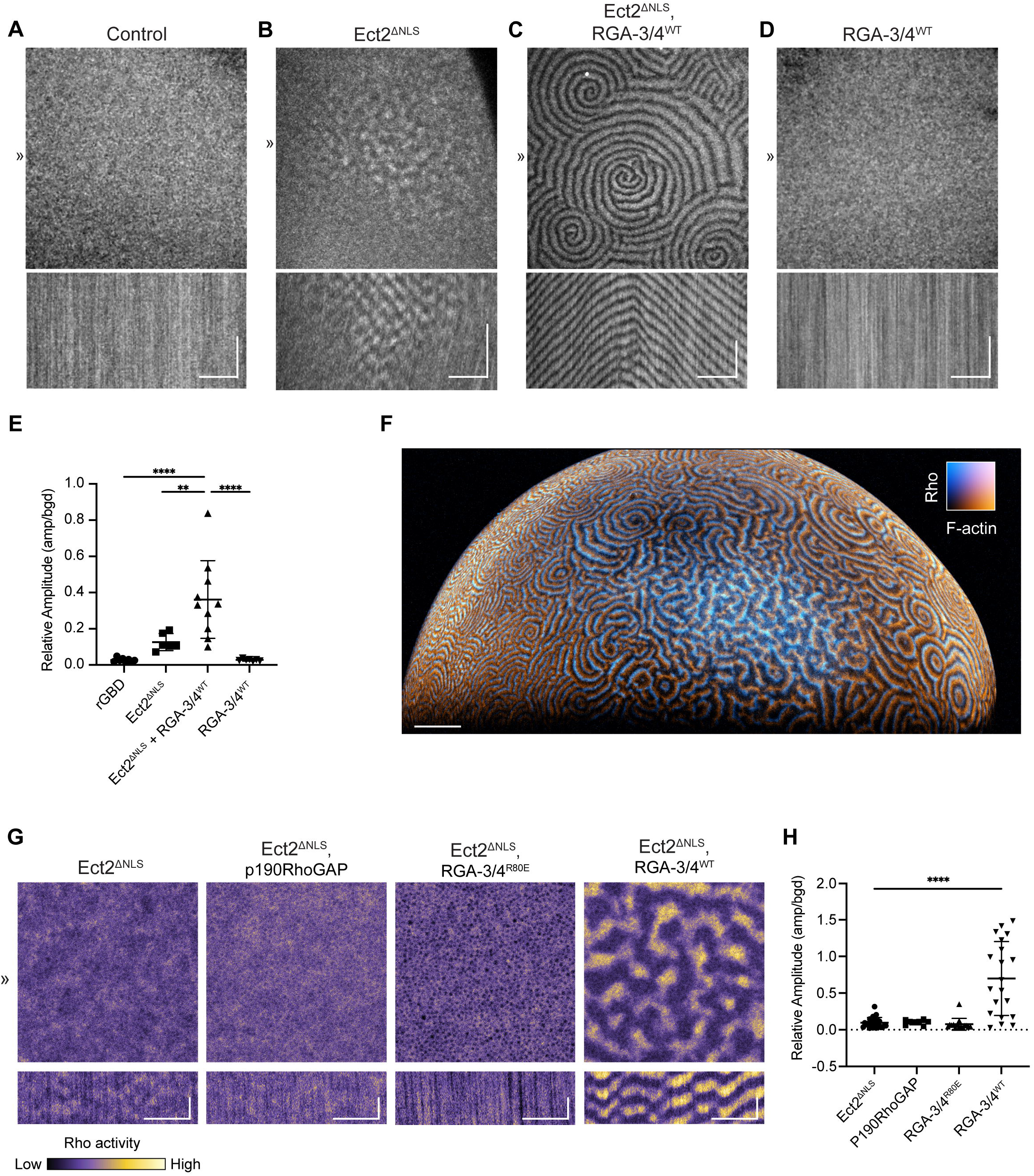
Co-expression of Ect2 and RGA-3/4 induces high-level excitability in immature frog oocytes. **(A-D)** Still frames (top) and kymographs (bottom) from representative oocytes expressing probe for active Rho (GFP-rGBD). Kymographs generated from 1px line drawn at position (»). x scale bar = 50 μm, y scale bar = 5 min; see also, Video 8. **(A)** Control oocyte expressing only active Rho probe shows no waves. **(B)** Oocyte expressing untagged, non-importable Ect2 (Ect2^ΔNLS^) shows isolated, low amplitude Rho waves. **(C)** Oocyte expressing Ect2^ΔNLS^ and RGA-3/4^WT^ shows high amplitude waves with multiple spiral cores and continuous waves across cortex. **(D)** Oocyte expressing RGA-3/4^WT^ shows no waves. **(E)** Quantification of relative wave amplitude across conditions A-D. Each dot represents a single oocyte; group mean ± standard deviation. One-way ANOVA with Tukey post-hoc test for multiple comparisons; **P<0.01; ****P<0.0001; rGBD, n = 8; Ect2^ΔNLS^, n = 6; Ect2 ^ΔNLS^ + RGA-3/4^WT^, n = 10; RGA-3/4^WT^, n = 7; 6 experiments. **(F)** Light-sheet imaging of immature oocyte expressing Ect2^ΔNLS^ and RGA-3/4^WT^ shows cortical waves present over entire animal cortex; scale bar = 100 μm; see also, Video 9. **(G)** All oocytes express probe for active Rho (GFP-rGBD) and Ect2^ΔNLS^. Expression of p190RhoGAP (panel 2) or RGA-3/4^R80E^ (panel 3) do not support high-level cortical excitability; x scale = 25 μm, y scale = 2 min. **(H)** Quantification of relative wave amplitude across experimental groups described in (G). Each dot represents a single oocyte; group mean ± standard deviation. One-way ANOVA with Tukey post-hoc test for multiple comparisons; ****P<0.0001; Ect2^ΔNLS^, n = 27; p190RhoGAP, n = 9; RGA-3/4^R80E^, n = 16; RGA-3/4^WT^, n = 21; 5 experiments.

The foregoing analysis indicated that the combination of Ect2^ΔNLS^ and RGA-3/4^WT^ induced cortical waves throughout the entire cortex. To better capture the scale of this phenomenon in these exceptionally large cells, we turned to light-sheet imaging, which permits rapid, relatively high-resolution imaging of large fields of view. Consistent with the impression obtained from imaging of smaller fields of view, light sheet imaging revealed that well-developed waves form throughout the oocyte cortex following coexpression of Ect2 and RGA-3/4^WT^ (Fig. 4F, Video 9). Further, this approach also permitted imaging for many hours with little loss of signal due to photobleaching.

Long-term imaging of cells co-expressing Ect2^ΔNLS^ and RGA-3/4 revealed that wave dynamics were well-developed at roughly 5 hrs post-injection and that same-day expression decreased cell-to-cell variability (not shown). Using this approach, we found that co-expression of Ect2^ΔNLS^ with the GAP-dead RGA-3/4 (RGA-3/4^R80E^) failed to induce cortical waves (Fig. 4G, H), demonstrating that GAP activity is needed to elicit this behavior. Co-expression of Ect2^ΔNLS^ with a different Rho GAP implicated in cytokinesis, p190RhoGAP (Manukyan et al., 2015; Su et al., 2003), also did not induce cortical waves in immature oocytes (Fig. 4G, H; Video 10), indicating that the effect of RGA-3/4 on cortical excitability is not replicated by an arbitrary choice of Rho GAP.

### Dynamic features of Rho, F-actin and RGA-3/4 in immature oocytes

The above results show that co-expression of Ect2 and RGA-3/4 generate complementary Rho and F-actin waves similar to those previously described in the cleavage furrows of starfish and frog embryos (Bement et al., 2015). To better understand the spatiotemporal relationship between RGA-3/4 and Rho we imaged tagged, wild-type RGA-3/4 (RGA-3/4^WT^-3xGFP) with a probe for active Rho (mCherry-rGBD) in the presence of Ect2^ΔNLS^. Consistent with the results obtained in starfish eggs, RGA-3/4^WT^-3xGFP was recruited to cortical waves that trailed Rho activity waves (Fig. 5A, B). Cross correlational analysis revealed that RGA-3/4^WT^-3xGFP signal peaked around 17.5 ± 3.7 s (n = 3 cells) after the peak of the signal for active Rho (Fig. 5C).

**Figure 5.**
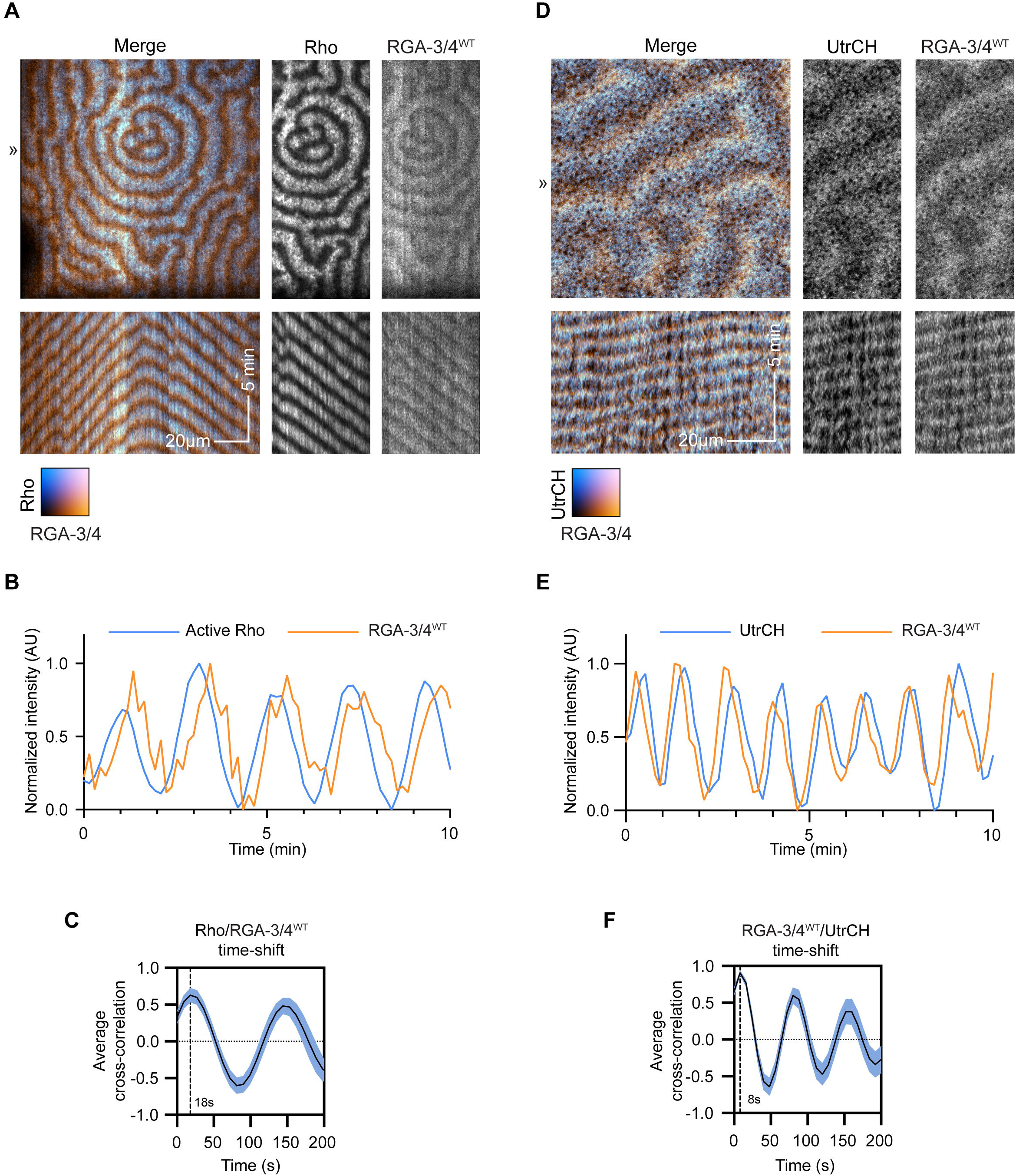
RGA-3/4 recruitment to waves trails Rho activation and slightly leads peak of F-actin recruitment. All oocytes expressing untagged Ect2^ΔNLS^ and RGA-3/4^WT^ to generate cortical waves. **(A)** Frog oocyte expressing probe for active Rho (cyan; GFP-rGBD) and tagged RGA-3/4^WT^ (orange; RGA-3/4^WT^-3xGFP). Kymographs (bottom) generated from 1px line drawn at position (»). **(B)** Representative intensity profile of active Rho and RGA-3/4^WT^. **(C)** Cross-correlational analysis of cell in (A) showing 18 s delay between Rho activation and RGA-3/4^WT^ recruitment. **(D)** Frog oocyte expressing probe for F-actin (cyan; mCherry-UtrCH) and tagged RGA-3/4 (orange; RGA-3/4^WT^-3xGFP). Kymographs (bottom) generated from 1px line drawn at position (»). **(E)** Representative intensity profile of F-actin and RGA-3/4^WT^. **(F)** Cross-correlational analysis of cell in (D) showing 8 s delay between peak RGA-3/4^WT^ recruitment and peak F-actin signal.

We then compared the dynamics of RGA-3/4 ^WT^-3xGFP to F-actin. As expected, based on the results obtained with starfish, there was significant overlap of RGA-3/4 ^WT^-3xGFP with mCherry-UtrCH (Fig. 5D, E). However, both qualitative assessment and cross-correlational analysis revealed that RGA-3/4 preferentially concentrated at the leading edge of the F-actin wave, with the peak actin signal trailing peak RGA-3/4 by 8.4 ± 2.4 s (n = 3 cells) (Fig. 5F).

### Recruitment of other cytokinetic participants to waves

If the Ect2 and RGA-3/4-generated waves in oocytes are indeed analogous to the waves found within the cleavage furrow, they should recruit a similar array of cytokinetic regulators. We therefore next assessed the behavior of known cytokinetic proteins in the immature oocyte system. Anillin—which binds Rho, F-actin, Ect2, and is commonly referred to as a “scaffolding protein” (Piekny and Glotzer, 2008; Piekny and Maddox, 2010)—was recruited to waves in immature oocytes, with peak intensity 18 s after the peak of active Rho (Fig. 6A-C). Myosin-2 waves were also observed trailing the peak of Rho activation, with a peak intensity roughly 57 s after the Rho peak (Fig. 6D-F); note that, because the F-actin assembly wave elicited by Rho lasts for a minute or so, this means myosin-2 largely associates with the trailing edge of the F-actin wave (in contrast to RGA-3/4 and anillin).

**Figure 6.**
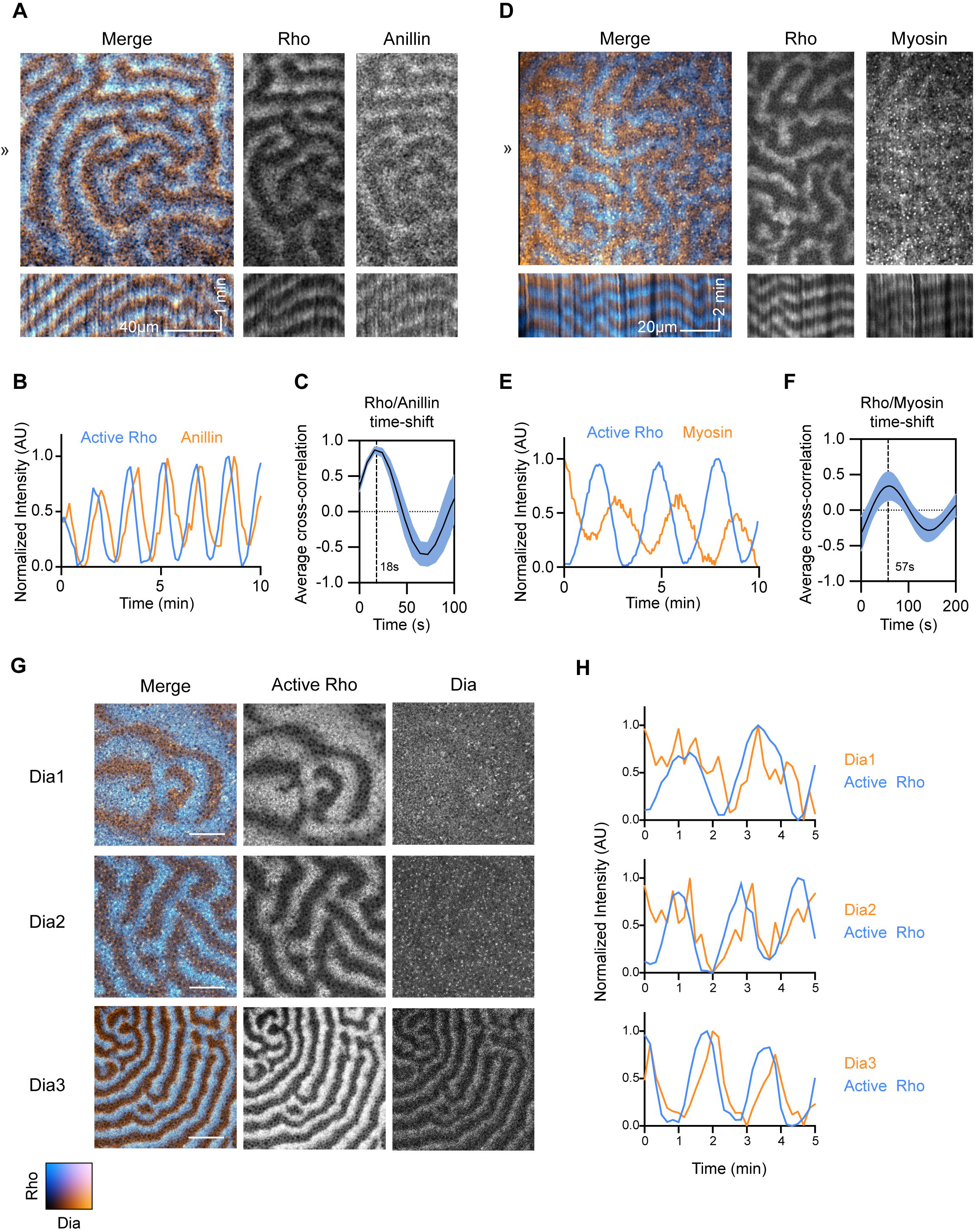
Recruitment of cytokinetic participants to immature oocyte waves. All oocytes expressing untagged Ect2^ΔNLS^ and RGA-3/4^WT^ to generate cortical waves. **(A)** Frog oocyte expressing probe for active Rho (cyan; mCherry-rGBD) and tagged Anillin (orange; Anillin-3xGFP). Kymographs (bottom) generated from 1px line drawn at position (»). **(B)** Representative Intensity profile of active Rho and Anillin. **(C)** Cross-correlational analysis of cell in (A) showing 18 s delay between Rho activation and Anillin recruitment. **(D)** Waving frog oocyte expressing probes for active Rho (cyan; mCherry-rGBD) and myosin (orange; Sf9-mNeon). Kymographs (bottom) generated from 1px line drawn at position (»). **(E)** Representative intensity profile of Rho and myosin dynamics for cell in (D). **(F)** Cross-correlational analysis of cell in (D) showing 57 s delay between Rho activation and Myosin recruitment. **(G)** Still frames of oocytes expressing probe for active Rho (cyan; mCherry-rGBD) and tagged *Xenopus* Dias 1, 2, or 3 (orange; Dia1-3xGFP, Dia2-3xGFP, Dia3-3xGFP). Only Dia3 is recruited robustly to cortical waves. Scale bar = 25 μm. **(H)** Representative intensity profiles of active Rho (cyan) with Dias 1, 2 and 3 (orange).

As a more stringent test of the cytokinetic “fidelity” of the immature oocyte system, we compared the behavior of the formins Dia1, Dia2 and Dia3, based on the previous demonstration that Dia3 is the major cytokinetic formin in *Xenopu*s embryos (Higashi et al., 2019). Dia2 showed very low-level recruitment to Rho waves in immature oocytes, while Dia1 showed essentially no recruitment (Fig. 6G, H). In contrast, Dia3 showed clear recruitment to the trailing edge of Rho waves (Fig. 6G, H). Thus, waves induced by Ect2 and RGA-3/4 in immature oocytes selectively recruit known participants of the cytokinetic apparatus assembly and function.

### Theoretical modeling of experiments with graded RGA-3/4 expression bridges dynamics in starfish and frog

Expression of RGA-3/4 strongly influences the wave characteristics in both activated starfish oocytes and immature frog oocytes, but in different ways. In starfish, overexpression of Ect2 alone induces dramatic high-amplitude waves, while increasing doses of RGA-3/4 reduced wave amplitude and width (Fig. 1; Fig. S1G, H). In frog, expression of RGA-3/4 and Ect2 were required for the emergence of high-amplitude waves with complex patterns (Fig. 4). This is surprising given that our results suggest that the F-actin-RGA-3/4 subsystem plays the role of the delayed negative feedback to Rho activation in both organisms. To gain insight into the observed behavior, we employed theoretical modeling of the Rho and F-actin dynamics.

Our model closely follows the approach we introduced previously (Bement et al., 2015). Briefly, the model (Fig. 7A) captures the salient processes leading to wave formation: (1) reversible membrane-cytoplasm shuttling of inactive Rho, (2) its Ect2-dependent positive-feedback activation, (3) F-actin polymerization stimulated by active Rho, (4) RGA-3/4-dependent Rho inactivation, and (5) the resultant F-actin disassembly. The detailed mathematical formulation of the model is given in the Methods. We chose the immature frog oocyte system as our experimental reference as it encompasses the broadest range of cortical behaviors and defined the model parameters accordingly, normalizing the fixed [Ect2] as unity and broadly varying [RGA-3/4] as a free parameter. This showed that at both very low and very high strengths of negative feedback, controlled by the value of [RGA-3/4], the model is found in stable spatially uniform states unable to support pattern formation. In agreement with intuition, the steady-state concentrations of active Rho and F-actin were higher at small [RGA-3/4] and lower at large [RGA-3/4]. We thus dubbed these states as the higher uniform state (HUS) and the lower uniform state (LUS), respectively (Fig. 7B). Linear stability analysis of the model showed that the two spatially uniform states are generically separated by the parameter domain of oscillatory behavior (green area, Fig. 7B), where the model is expected to exhibit a variety of dynamic wave patterns. Furthermore, we found that waves should be also observed in the parameter regions (blue area, Fig. 7B) in which the spatially homogeneous state is destabilized by diffusion. This is caused by the oscillatory analogue of the Turing mechanism known as *wave instability* (Vanag and Epstein, 2009). Thus, as [RGA-3/4] is increased, waves are expected to spontaneously appear at the boundary between the domains of HUS and wave instability and persist throughout the wave instability and oscillation domains (Fig. 7B).

**Figure 7.**
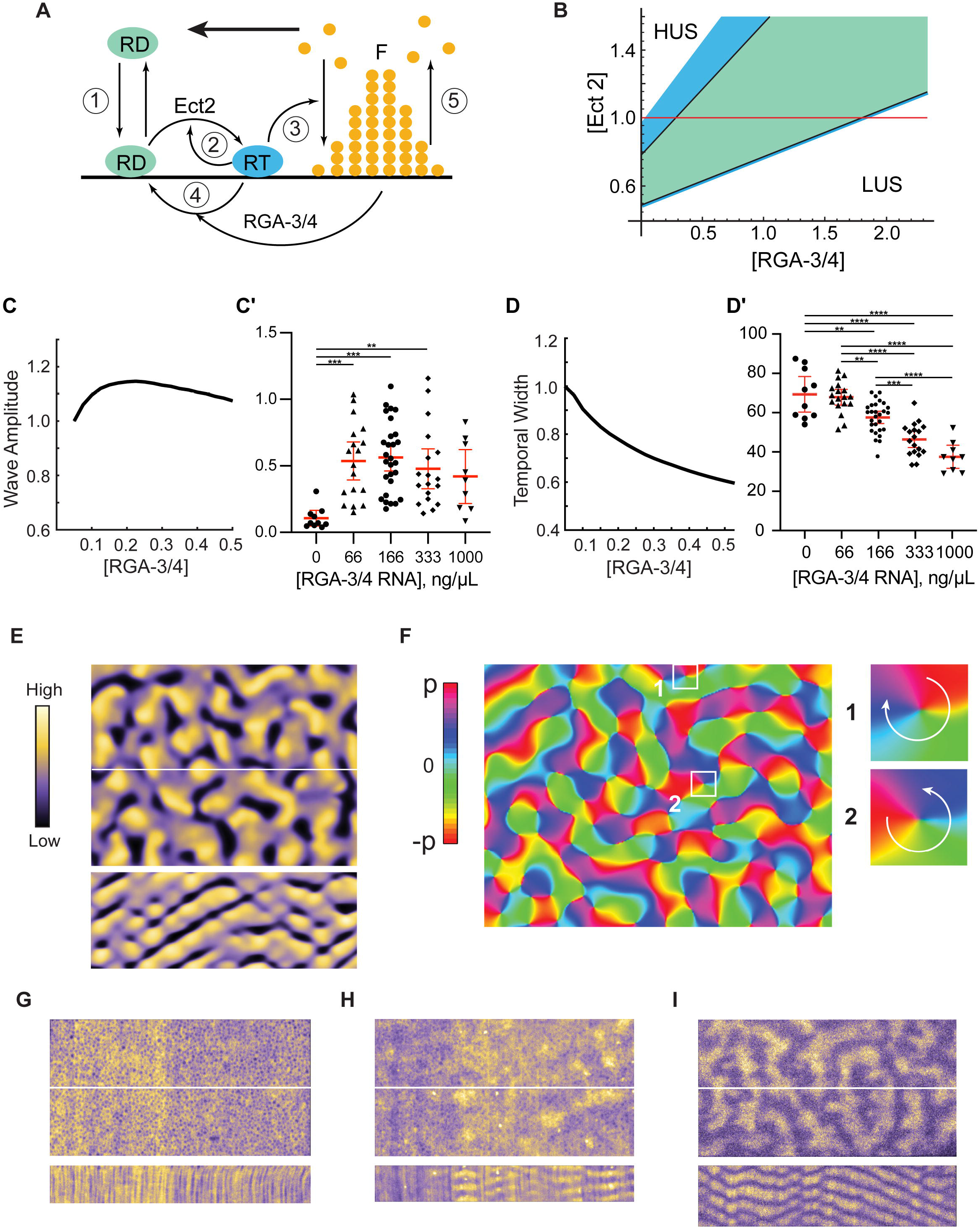
The model robustly predicts wave dynamics preceded by a turbulence regime. **(A)** The major reactions described by the model (see text and Methods for details). RD – inactive Rho; RT – active Rho; F – F-actin. Bold arrow indicates the direction of wave propagation. **(B)** Diagram of the model behavior, waves are predicted in the domains of wave instability (blue) and oscillations (green). HUS, LUS – higher and lower uniform states. **(C-C’)** Modeling (C) vs in vivo (C’) data of normalized active Rho wave amplitude over changing [RGA-3/4^WT^]. **(D-D’)** Modeling (D) vs in vivo (D’) data of normalized Rho wave temporal width over changing [RGA-3/4 ^WT^]. In (C’, D’), each dot represents a single oocyte; group mean ± 95% CI; 0 ng/μL, n = 10; 66 ng/μL, n = 18; 166 ng/μL, n = 28; 333 ng/μL, n = 18; 1000 ng/μL, n = 9; 7 experiments. One-way ANOVA with Tukey post-hoc test for multiple comparisons; **P<0.01; ***P<0.001; ****P<0.0001. **(E)** Spiral turbulence induced by noise on the boundary between the HUS and the wave instability domain in the model. Active Rho amplitude is color-coded (left). Still-frame (top) and kymograph (bottom); kymograph computed from the central white line. **(F)** The computationally reconstructed phase of the wave dynamics in (E). Turbulent behavior is induced by formation and motion of pairs of phase defects with the opposite charge. A representative pair of defects is shown in insets. Phase increases clockwise in 1 (charge +1), while counterclockwise in 2 (−1). **(G-I)** Still-frames of color-coded Rho activity (top) and resulting kymographs (bottom). (G) Rho flickers at 0 ng/μL. (H) Turbulence 1 at 33 ng/μL. (I) Fully developed turbulence 2 at 66 ng/μL.

To test these predictions, we performed detailed model simulations, which produced wave patterns precisely within the parameter range predicted by the stability analysis. In the model, as RGA-3/4 was increased from 0, waves emerged with a finite amplitude that reached a maximum at the boundary between the wave instability and oscillatory domains and then diminished (Fig. 7C). The wave amplitude measured in experiments showed the same trend, with even low levels of exogenous RGA-3/4 resulting in a sharp increase in wave amplitude (Fig. 7C’). In a striking quantitative agreement, both the model and experiment found that the temporal width of the waves monotonically decreased with the increase in [RGA-3/4] (Fig. 7D, D’). Importantly, the same behavior was also observed in the activated starfish oocytes (Fig. S1G, H). The time period of waves also diminished with [RGA-3/4] both in the model and in experiment (Fig. S4A, A’).

Remarkably, molecular noise incorporated into our model brought out behavior that could not be predicted by a purely deterministic model or gleaned from the stability analysis. On the boundary between the HUS and the domain of wave instability, the model exhibited highly irregular, rapidly changing patterns of activity characterized by chaotically moving wave fragments. At small [RGA-3/4], they emerged as infrequent Rho activity pulses that propagated only a short distance prior to disappearing. As [RGA-3/4] increased, these spatially isolated pulses coalesced into the progressively longer wave fragments that occupied the entire spatial domain, completely replacing the HUS (Fig. 7E, top). Despite a chaotic spatial appearance, kymographs of these dynamics revealed a remarkable periodicity in time (Fig. 7E, bottom). Numeric reconstruction of the oscillation phase of these periodic dynamics (Bement et al., 2015) showed that it is dominated by the erratic motion of phase defects (Fig. 7F). These defects (Winfree, 1980), which also serve as the cores of spiral waves, always emerge and disappear in pairs with the opposite charge (Fig. 7F, insets 1&2). Their random pair-wise creation and annihilation interspersed by the intervals of irregular motion create an exotic condition known as *spiral turbulence* (Aranson and Kramer, 2002).

Comparison of our model predictions with the experimental observations revealed both an astonishingly rich diversity of cortical behaviors and surprising qualitative similarity between the model and experimental results (Fig. 7E, G-I, Fig. S4B). In the absence of exogenous RGA-3/4, cells predominantly showed localized flickers of Rho activity that did not exhibit any propagation (Fig. 7G). This heightened Rho activity likely corresponds to the HUS in the model. A smaller proportion of samples exhibited erratically moving isolated maxima of Rho activity (Fig. 7H, Videos 8, 10), reminiscent of the pulses in the model. To distinguish this spatiotemporal dynamical regime from the fully developed spiral turbulence, or turbulence 2 (T2), we dubbed it turbulence 1 (T1). The addition of even low levels of exogenous RGA-3/4 resulted in the replacement of static Rho flickers by T1 and T2 dynamics. In the regime of the fully developed spiral turbulence (T2, Fig. 7I, Video 10), readily recognizable propagating wave fragments densely populated extended fields of view.

Modeling showed that further increase in [RGA-3/4] induced a rapid transition from the spiral turbulence to the periodic propagation of wave trains. While we observed a continuous change in the wave patterns, they could nevertheless be partitioned approximately into two classes. The first class, typical of the wave instability domain, was represented by wave trains of thick, gently curved waves (Fig. 8A). The second class, characteristic of the oscillatory domain, was represented by thin waves that formed multiple involute spirals typically showing only a fraction of the turn (Fig. 8B). Patterns falling within each class could be readily identified in the experimental results (see Figs. 8A’, B’ Video 11). The model and experimental wave patterns shared unique morphological signatures formed by the co-occurrence of characteristic features, such as dislocations (arrowheads, Figs. 8A, A’), grain boundaries (Fig. 8C), and two-armed spirals (Fig. 8D). A peculiar type of line defect (Goryachev et al., 2000; la Porta and Surko, 1996) was observed in both the model and experiment at the interface of two wave trains whose wave vectors are nearly antiparallel (yellow dashed line, Figs. 8B, B’, E). Fig. 8F summarizes the relative frequency of ten morphological features observed in the immature frog oocyte injected with the increasing concentrations of RGA-3/4 (see Methods for details). As in the model, spatiotemporal dynamics in vivo rapidly progressed from the spatially uniform state with high Rho activity (no exogenous RGA-3/4) via the characteristic succession of chaotic turbulent states, to the fully developed wave patterns, first with the signature of wave instability and then that of the oscillatory regime (Fig. 8F; Fig. S4B).

**Figure 8.**
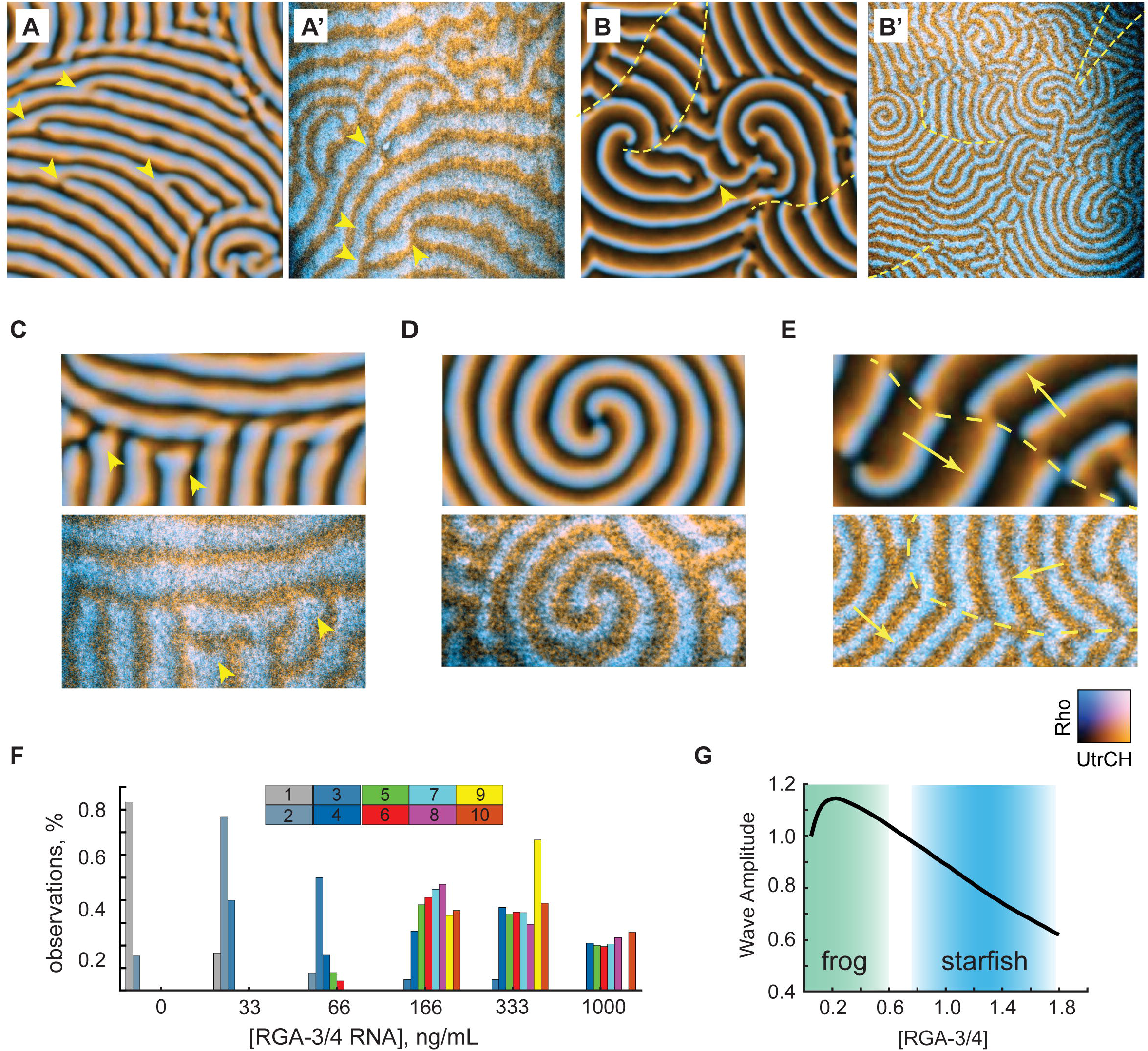
The model reproduces qualitative morphological signatures of experimental wave patterns. (A) Model wave pattern typical of the wave instability domain. (A’) Representative experimental wave pattern. Arrowheads in (A, A’) point to wave front dislocations. See also, Video 11. (B) Model wave pattern typical of the oscillatory domain (B’) Representative experimental wave pattern. Arrowhead in (B) points to dislocation. Yellow dash lines in (B, B’) mark line defects separating wave trains (see E). See also, Video 11. **(C-E)** Characteristic wave pattern features in the model (top) and experiment (bottom). (C) A grain boundary; arrowheads point to the wave front ends with a typical bulbous morphology. (D) Stable two-armed spirals with multiple turns. (E) A line defect (dash line) on the interface of two wave trains whose wave vectors are shown by yellow arrows. **(F)** Relative observation frequency of ten morphological features of wave patterns (see Methods for details). (G) Biphasic behavior of the wave amplitude in the model and the qualitative mapping of the two experimental systems onto the model. Color transparency indicates confidence of mapping boundaries.

As the strength of negative feedback is increased in the model, the system crosses the parameter domain of wave dynamics, whose amplitude shows asymmetric biphasic behavior, first rapidly increasing and then slowly diminishing until “crashing” onto the lower uniform state (Fig. 8G). Using the combination of quantitative measures, such as the wave amplitude and temporal width, and morphological features of wave pattern, we can confidently map the immature frog oocyte system onto the lower end of the negative feedback strength in the model (green area, Fig. 8G). By the same argument, activated starfish oocyte system maps on the higher end of the negative feedback strength in the model (blue area, Fig. 8G). The model thus effectively bridges the two experimental systems and indicates that their “ground levels” of negative feedback strength may be substantially different.

## Discussion

This study demonstrates that Rho, Ect2, RGA-3/4 and F-actin form the core of a circuit that regulates cytokinesis and cortical dynamics in both starfish and frogs. This conclusion is based on these findings: First, RGA-3/4 localizes to the equatorial cortex prior to and during cytokinesis. Second, RGA-3/4 waves “chase” Ect2-induced Rho activity waves, with RGA-3/4 recruitment closely matching Rho-induced F-actin assembly waves. Third, experimental elevation of RGA-3/4 narrows and lowers Rho waves in a GAP-dependent manner. Fourth, latrunculin-induced changes in F-actin wave organization produces corresponding changes in the RGA-3/4 waves. Fifth (and most compellingly), the co-expression of RGA-3/4 and Ect2 is sufficient to induce complex, high-amplitude, and frankly psychedelic waves of active Rho, F-actin, and cytokinetic regulators in the otherwise inert cortex of the immature oocytes. Thus, an image emerges in which cell fission results from at least two, coupled, Rho-dependent feedback loops focused at the equatorial cortex: a positive feedback loop mediated by Ect2 and a delayed negative feedback loop acting through RGA-3/4 and F-actin.

Are these results specific to frogs and starfish? Likely not: previous studies of RGA-3/4 in HeLa cells and *C. elegans* embryos found that RGA-3/4 negatively regulates Rho during cytokinesis, as judged by increased contractility and recruitment of the Rho targets anillin and Rho-dependent kinase (Bell et al., 2020; Zanin et al., 2013). Further, RGA-3/4 was found to antagonize Rho activity in an F-actin dependent manner during cortical flow and pulsed contraction in *C. elegans* (Michaux et al., 2018), indicating that in this system too, F-actin and RGA-3/4 collaborate to produce cortical excitability. Collectively, these results indicate that Ect2, RGA-3/4, F-actin and Rho represent part of a conserved cortical circuit with a repertoire that spans focal contraction, steady wave propagation, and spatial patterning at varied scales.

One of the most exciting discoveries of this study is that the cortex of immature oocytes can be transformed from an inert quiescent state to a highly active state by the expression of just two proteins. The oocyte system overcomes several limitations of the natural excitability evident in embryos. First, immature oocytes are arrested in an interphase-like state and thus are not subject to the rapid cell-cycle and developmental changes intrinsic to early embryos, greatly reducing cell-to-cell variation and analysis complexity (Bement et al., 2015; Swider et al., 2022). Second, capturing cortical waves in the fleeting embryonic furrow is extremely challenging, especially as cells become smaller. In contrast, the oocyte system provides a large-scale, quasi-two dimensional representation of the furrow, which, combined with the persistence of the waves, makes it much easier to acquire high resolution data. Finally, precisely because the immature oocyte is not immediately prepared for division or motility or any other large-scale shape change, the influence of different cytokinetic participants (or other factors) on cortical dynamics can be systematically and quantitatively assessed simply by microinjection of varying concentrations of mRNA. The power of this approach is illustrated by the qualitative and quantitative variation in cortical dynamics observed by graded RGA-3/4 expression. Further, given that the system possesses cytokinetic “fidelity”—in that known cytokinetic targets of Rho are also recruited to waves—we anticipate that it will prove a highly useful counterpart to more traditional approaches used to study conserved cytokinetic regulators thought to impact Rho dynamics.

The strengths of the immature oocyte system will also prove useful for studies beyond cytokinesis or even biology, as robust experimental systems exhibiting the complex spatiotemporal dynamics it produces are in short supply. Principle among these are the classic Belousov-Zhabotinsky (BZ) reaction (Zaikin and Zhabotinsky, 1970) and the more-recently developed bacterial MinD system reconstituted on supported lipid bilayers (Brauns et al., 2021; Loose et al., 2008), each of which can, with the appropriate manipulations, produce a diversity of excitable and oscillatory patterns. As the oocyte represents a living counterpart to these in vitro systems that rivals them in pattern complexity, it is likely that it will be of interest to those in the fields of mathematics and physics who specialize in the study of complex self-organized patterns.

Just how versatile is the pattern forming circuit described here? Simply varying the level of exogenous RGA-3/4 produces a broad range of dynamic patterns, ranging from Rho pulses, to short, “choppy” waves, to longer, labyrinth-like waves, to fully developed spiral waves. When one considers other aspects of the circuit, the potential for further pattern variation becomes enormous. For example, a recent study demonstrated that suppressing expression of specific Rho effectors induces transition from excitatory pulses to noisy oscillatory pulses or noisy oscillatory waves (Yao et al., 2022). Our modeling results in the current study further underscore the potential for novel patterns as the starfish system differs from the immature oocyte system in that it apparently operates in a domain with higher basal negative feedback, implying that if this situation could be mimicked in the immature frog oocyte system, yet further patterns could be generated. While the circuit versatility is particularly evident in the immature oocyte model, we suspect other cortical circuits will likewise prove versatile, as suggested by a recent study of cortical dynamics in Dictyostelium (Yochelis et al., 2022).

The modeling results make two additional, and important points: We and others have reported experimental observation of spiral turbulence in starfish oocytes (Bement et al., 2015; Tan et al., 2020) and activated frog eggs overexpressing Ect2 (Bement et al., 2015). Earlier work studied spiral turbulence in a non-system-specific abstract model (Aranson and Kramer, 2002). Here, we demonstrate that the introduction of noise is sufficient to induce spiral turbulence in a biologically realistic model of the cellular cortical dynamics. Second, the extensive similarity between the model and experimental results suggests that even a simple biophysical model lacking fine biochemical details can predict how the spatiotemporal dynamics of a complex in vivo system will change with variation in the strength of positive and negative feedback.

One of the most important results to arise from the manipulation of negative feedback strength is that elevating expression of RGA-3/4—a Rho inactivator—sharply *increases* Rho wave amplitude such that the cell focusses relatively more Rho activity in specific areas of the cortex (i.e. at the peaks of the Rho waves). This finding is consistent with earlier studies proposing the existence of GAP-driven Rho GTPase turnover flux as a means to counter the effects of diffusion (Bement et al., 2006; Goryachev and Pokhilko, 2006). While evidence for rapid Rho GTPase turnover has since been provided in several studies (e.g., Miller and Bement, 2009; Burkel et al., 2012; Budnar et al., 2019), the results reported here directly demonstrate Rho GAP-induced elevation of local Rho activity. This seemingly paradoxical result is in fact consistent with the ability of GAPs to reduce GTPase activity. That is, although addition of RGA-3/4 increases wave amplitude, the *total* amount of Rho activity diminishes as RGA-3/4 concentration is increased.

One final puzzle must be considered: while we are persuaded that RGA-3/4 directly or indirectly associates with F-actin based both on the work here and that of Michaux et al. (2018), the fact that the peak of the RGA-3/4 waves slightly leads the peak of the F-actin waves in both the frog and starfish system suggests that the association may be more complex than direct binding of RGA-3/4 to bulk F-actin. Several nonexclusive possibilities suggest themselves: perhaps RGA-3/4 preferentially associates with a particular pool of F-actin that only represents a part of the total F-actin comprising the waves, as observed in yeast, which recruit distinct actin-binding proteins to Arp2/3-versus formin-nucleated filaments (Kadzik et al., 2020). It is also possible that RGA-3/4 associates with F-actin indirectly, as suggested by the recent demonstration that its recruitment to cortical foci involved in pulsed contractions in *C. elegans* is dependent on two proteins recently identified as cytokinesis regulators (Bell et al., 2020). Finally, while the primary structure of RGA-3/4 is poorly conserved across species, structural predictions indicate that much of the protein is intrinsically disordered (Jumper et al., 2021), so it may be that its recruitment to F-actin promotes a liquid-liquid phase transition of RGA-3/4, analogous to the situation for TPX2 recruitment to microtubules (King and Petry, 2020). In any case, further work aimed at identifying RGA-3/4’s mode of recruitment, as well as its interaction profile, will greatly benefit the field of cortical excitability and cytokinetic signaling.

---

**Supplemental Figure 1.**
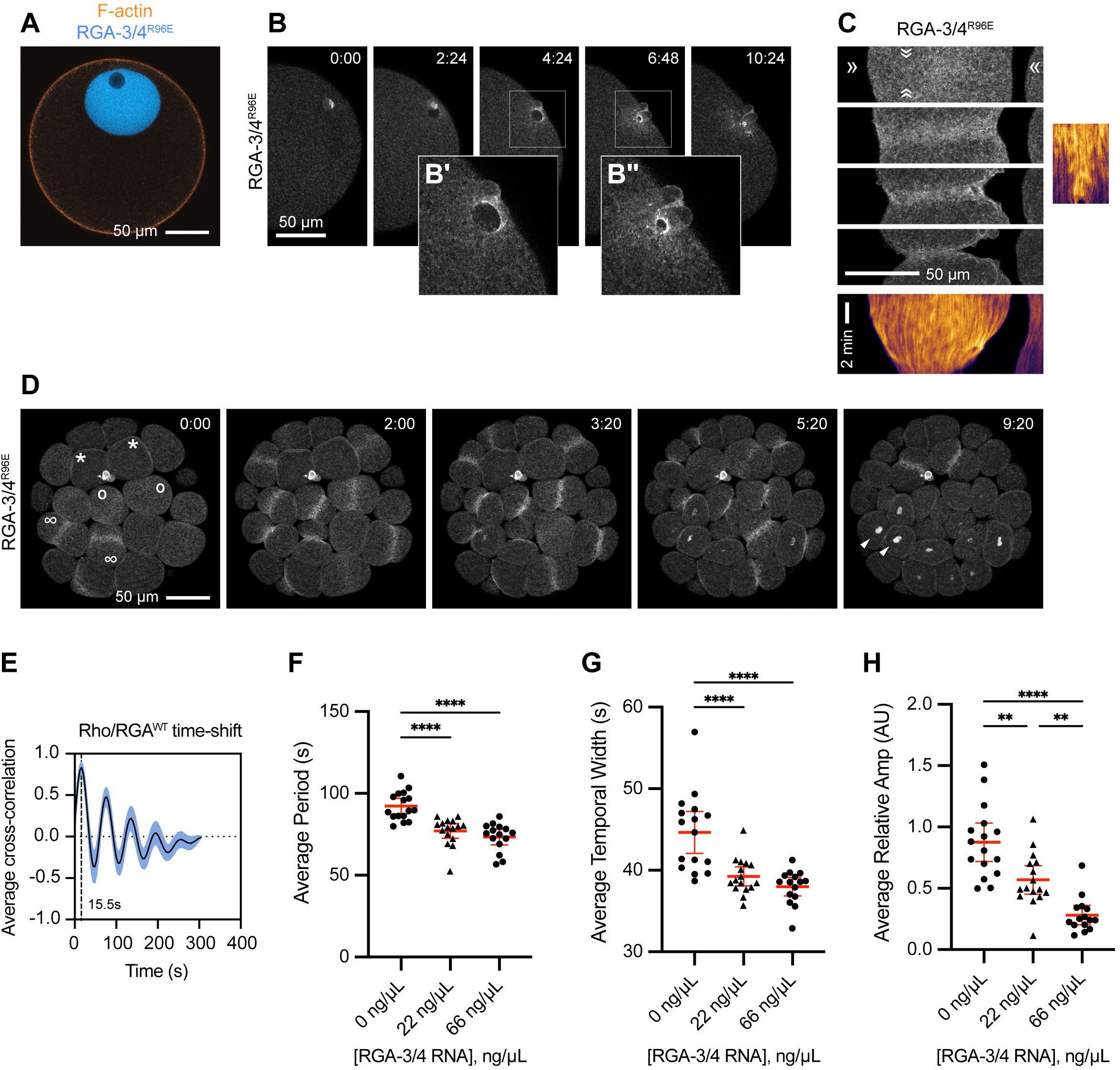
Localization of RGA-3/4 to the germinal vesicle, meiotic cytokinetic apparatus, nucleus, and mitotic cytokinetic apparatus in starfish. (**A**) Localization of GAP-dead RGA-3/4 (mNeon-RGA-3M^R96E^; cyan) and F-actin (mCherry-UtrCH; orange) to the germinal vesicle and cortex of the immature starfish oocyte, respectively. (**B**) Time course of mNeon-RGA-3/4^R96E^ localization during second meiosis. Time in min:sec. Insets **B’, B’’** enlarge nascent cytokinetic apparatus 2.5x as indicated by boxes **(C)** mNeon-RGA-3/4^R96E^ recruitment to the equatorial cortex during cytokinesis and corresponding kymographs – note low-amplitude wavelets throughout furrow ingression. **(D)** mNeon-RGA-3/4^R96E^ localization in cleaving blastomeres of 32-cell starfish embryo; asterisks indicate cells in interphase; o’s indicates cells in early anaphase – note cortical accumulation of mNeon-RGA-3/4^R96E^ compared to interphase cells; ∞’s indicate cells that have commenced cytokinesis; arrowheads indicate reforming nuclei. Time in min:sec. **(E)** Cross-correlational analysis of a starfish cell expressing mNeon-RGA-3/4^WT^ and mCherry-rGBD showing a 15 s delay between peak Rho activity and RGA-3/4^WT^ recruitment. Corresponds to experiments shown in Fig. 1C (75ng/μl). **(F-H)** Quantification of period (F), temporal width (G) and relative amplitude (H) for experiments shown in Fig. 1E. Each dot represents a single oocyte; group mean ± 95% CI; 0 ng/μL, n = 16; 22 ng/μL, n = 16; 66 ng/μL, n = 15; 2 experiments. One-way ANOVA with Tukey post-hoc test for multiple comparisons; **P<0.01; ****P<0.0001.

**Supplemental Figure 2.**
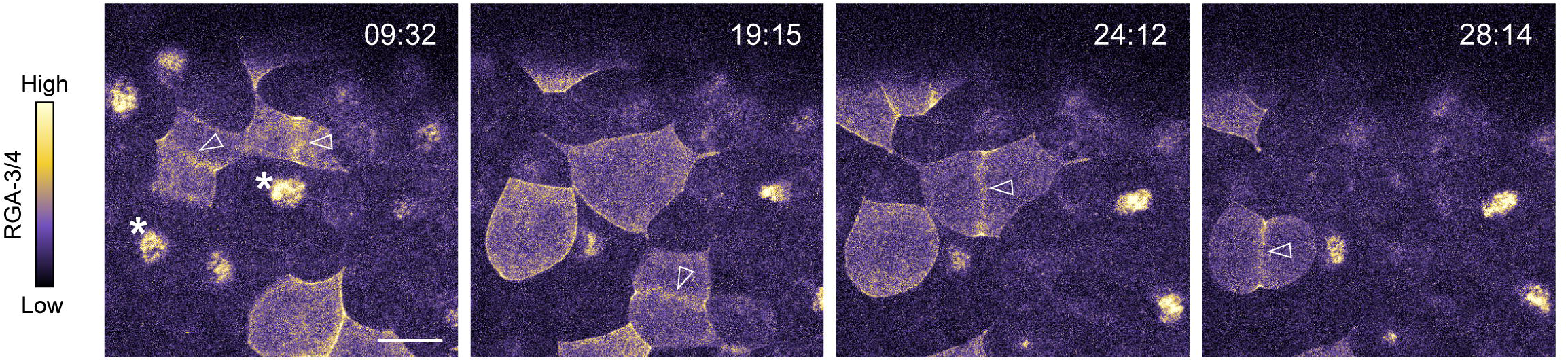
Time course of a frog embryo expressing RGA-3/4 ^WT^-3xGFP. Time in min:sec since start of recording. RGA localizes to equatorial cortex and contractile ring (white arrow heads) in cells undergoing cytokinesis and nuclei (white asterisks) in interphase cells. Scale bar = 25 μm. See also, Video 5.

**Supplemental Figure 3.**
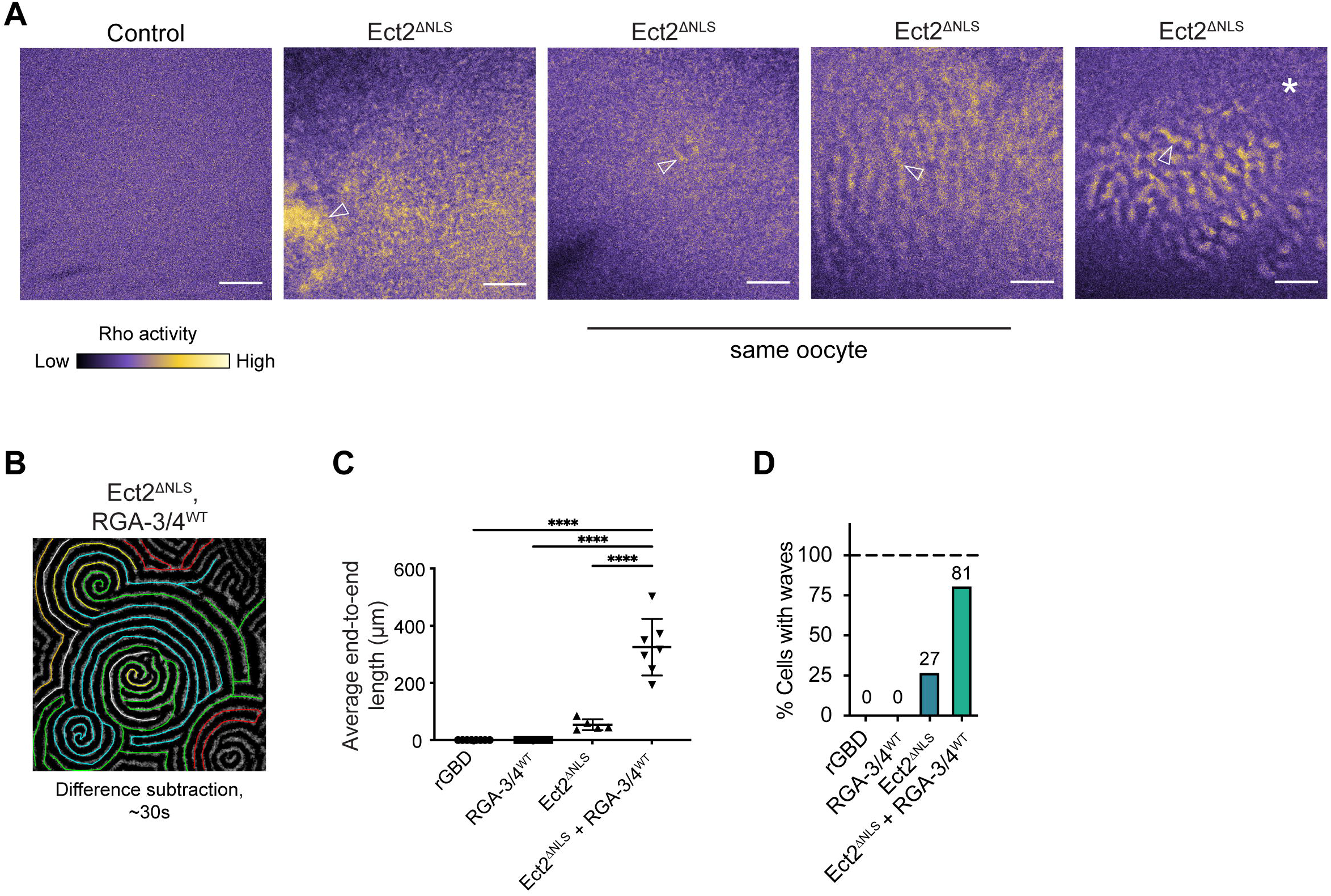
Variation in Rho activity patterns in oocytes expressing Ect2 alone and quantification for oocytes co-expressing Ect2 and RGA. **(A)** All oocytes express probe for active Rho (GFP-rGBD). Panel 1, rGBD only; panels 2-5, examples of phenotypes from Ect2^ΔNLS^ overexpression: static patches of Rho activity but no traveling waves (panel 2, arrowhead); tiny cluster of waves (panel 3, arrowhead) and diffuse wave patterns (panel 4, arrowhead) in same oocyte; wave patches (panel 5, arrowhead), surrounded by dormant cortex (panel 5, asterisk); scale bars = 50 μm. **(B)** Example still frame difference subtraction of oocyte from (Fig. 4C), showing segmentation process for measuring end-to-end lengths of cortical waves. **(C)** One-way ANOVA with Tukey post-hoc test for multiple comparisons, comparing end-to-end lengths across experimental groups. Each dot represents a single oocyte; group mean ± standard deviation. Cells co-expressing Ect2^ΔNLS^ and RGA-3/4^WT^ are significantly different from all other groups; controls, n = 8; RGA-3/4^WT^, n = 7; Ect2^ΔNLS^, n = 11; Ect2^ΔNLS^ + RGA-3/4^WT^, n = 12; 7 experiments; ****P<0.0001. **(D)** Plot of percent of cells displaying cortical waves across each experimental condition; controls, n = 8; RGA-3/4^WT^, n = 7; Ect2^ΔNLS^, n = 36; Ect2^ΔNLS^ + RGA-3/4^WT^, n = 37; 13 experiments.

**Supplemental Figure 4.**
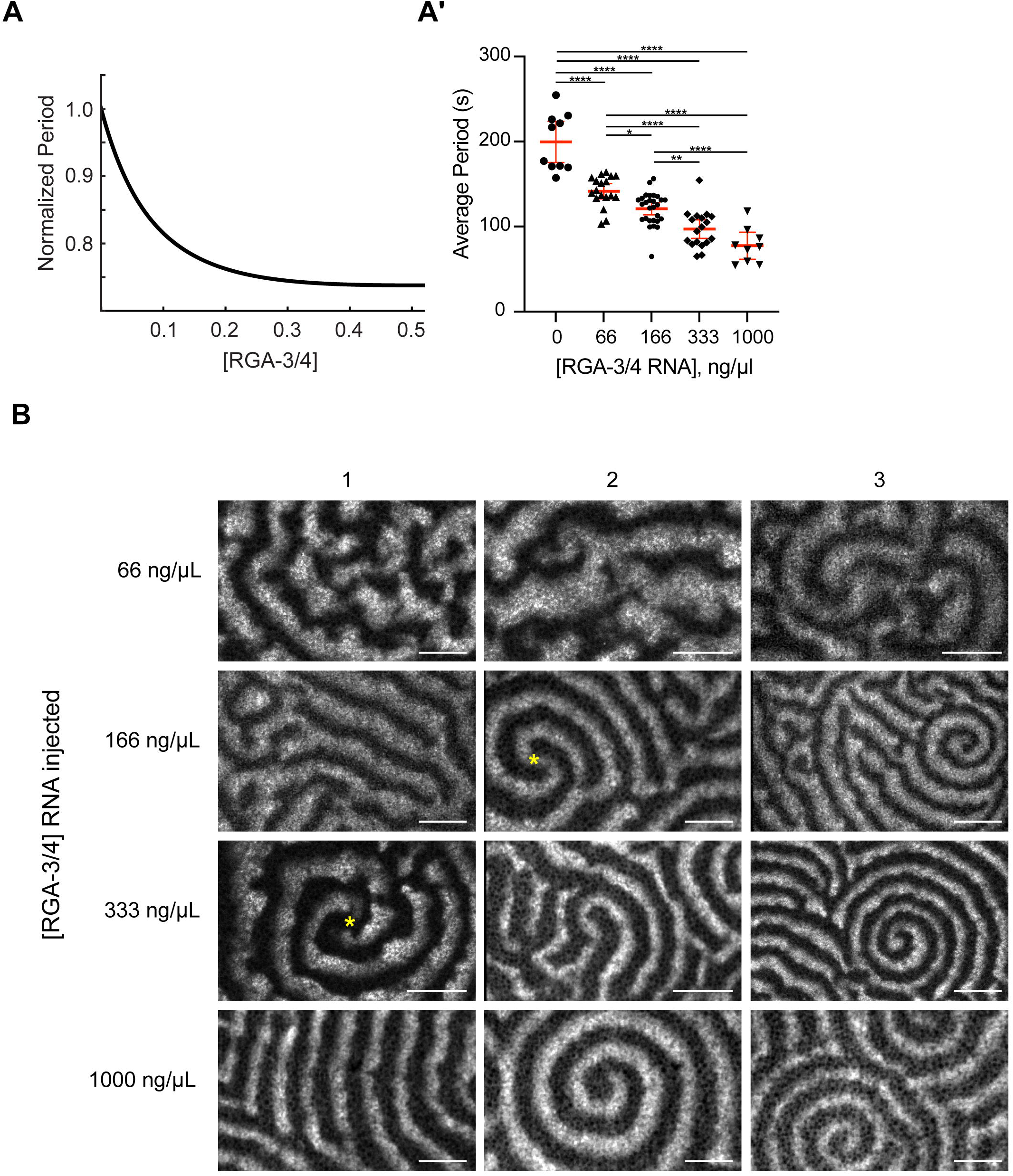
Changes in the activator:inhibitor ratio produce a wide range of cortical behaviors. (A-A’) Modeling (A) and in vivo (A’) data of normalized Rho wave period over changing [RGA-3/4^WT^]. Each dot represents a single oocyte; group mean ± 95% CI; 0 ng/μL, n = 10; 66 ng/μL, n = 18; 166 ng/μL, n = 28; 333 ng/μL, n = 18; 1000 ng/μL, n = 9; 7 experiments. One-way ANOVA with Tukey post-hoc test for multiple comparisons; *P<0.05; **P<0.01; ***P<0.001; ****P<0.0001. **(B)** Representative oocytes from the quantifications shown in Fig. 7C’, D’ and Fig. S4A’. Each row represents 3 individual cells at the noted [RGA-3/4^WT^]. Waves progress from choppy/turbulent spirals to long unbroken spiral wave chains that dominate the cortex. Scale bar = 25 μm. Yellow asterisks represent double spiral cores.

**Video 1.** Post-meiotic starfish oocytes expressing mCh-rGBD (cyan; right), excess wildtype Ect2, and varying doses of mNeon-RGA-3/4^WT^ (orange; left): 25 ng/μl (needle concentration), 75 ng/μl, and 200 ng/μl. Corresponds to Figure 1C. Time in min:sec from start of recording; all are single superficial optical planes at 4 sec intervals. Sequences highlight two key points: 1) RGA-3/4 recruits in the wake of Rho activity waves, and 2) increasing dose of RGA-3/4 modulates waves, first regularizing them before suppressing their amplitude and continuity.

**Video 2.** Post-meiotic starfish oocyte expressing mCh-rGBD (cyan; right), excess wildtype Ect2, and 250 ng/μl (needle concentration) mNeon-RGA-3/4^R96E^ (orange; left). Corresponds to Figure 1D. Time in min:sec from start of recording; single superficial plane at 4 sec intervals. Sequence demonstrates the phase relationship between Rho activity and RGA-3/4 recruitment: throughout the field, orange follows blue closely. Dose of Ect2 titrated to evoke regular rolling waves of Rho activity; R96E mutant has no detectable effect on waves.

**Video 3.** Post-meiotic starfish oocytes expressing excess wild-type Ect2, GFP-rGBD (cyan; right), mCh-UtrCH (orange; left), and varying doses of untagged wild-type starfish RGA-3/4: 0, 22 ng/μl, and 66 ng/μl. Corresponds to Figure 1E. Time in min:sec from start of recording; all are maximum projections of 3 superficial optical planes at 8 sec. intervals. Sequences illustrate the progressive effect of increasing RGA-3/4 dose on Rho wave amplitude and behavior: sloppy high-amplitude bursts are converted to orderly high-amplitude propagating fronts, then to low-amplitude broken fronts with limited propagation.

**Video 4.** Post-meiotic starfish oocyte expressing excess wild-type Ect2, GFP-rGBD (cyan; left), and mCh-UtrCH (orange; right), treated at time 0 with 200 nM Latrunculin B. Corresponds to Figure 2B. Time in min:sec relative to time of Latrunculin perfusion. Latrunculin remains present for the duration of the sequence, which shows that this treatment noticeably increases Rho wave amplitude and duration.

**Video 5.** Time course of a late-stage frog embryo expressing RGA-3/4 ^WT^-3xGFP. Corresponds to Figure S2. Time in min:sec since start of recording; 11 seconds/frame at 20 fps. RGA localizes to equatorial cortex and contractile ring in cells undergoing cytokinesis and nuclei in interphase cells.

**Video 6.** Frog embryo expressing RGA-3/4 ^WT^-3xGFP (orange) and a probe for F-actin (UtrCH; cyan) undergoing cytokinesis. Corresponds to Figure 3A. 11 seconds/frame at 20 fps. RGA-3/4 signal localizes to the cortex upon nuclear envelope breakdown and colocalizes at the contractile ring with F-actin during cytokinesis.

**Video 7.** Localization of GAP-dead RGA-3/4 (RGA-3/4^R80E^-3xGFP) in an early frog embryo. Corresponds to Figure 3B. Time in min:sec since start of recording; 4 seconds/frame at 20 fps. Waves of RGA-3/4 activity can be seen outside and inside the furrow.

**Video 8.** Oocytes expressing a probe for active Rho (rGBD) and various combinations of Ect2^ΔNLS^ and RGA-3/4^WT^. Corresponds to Figure 4A-D. Time in min:sec since start of recording; 9, 7, 10, and 9 seconds/frame (Fig 4.A-D) at 20 fps. Cells co-expressing Ect2^ΔNLS^ and RGA-3/4^WT^ show marked increase in excitable dynamics.

**Video 9.** Light-sheet movie of an oocyte co-expressing untagged Ect2^ΔNLS^ and wild-type RGA-3/4, along with probes for active Rho (cyan) and F-actin (orange). Corresponds to Figure 4F. Time in hh:mm:ss; 25 seconds/frame at 20fps. Oocyte is oriented with animal hemisphere enface and half of the hemisphere was captured.

**Video 10.** Video corresponding to Figure 4G. Oocytes expressing a probe for active Rho (rGBD) and combinations of Ect2^ΔNLS^ and either RGA-3/4^WT^, RGA-3/4^R80E^ or p190RhoGAP. Only cells with wild-type RGA-3/4 display high-level cortical excitability. Time in min:sec since start of recording; 6 seconds/frame at 20 fps.

**Video 11.** Examples of agreement between computational simulations and in vivo data. Simulations show active Rho (cyan) and F-actin (orange) waves. In vivo data is from oocytes expressing untagged Ect2^ΔNLS^ and RGA-3/4 mRNA, along with probes for active Rho (cyan) and F-actin (orange). Corresponds to Figure 8A-B’.

## Methods

### Starfish oocytes

*Patiria miniata* were obtained from Marinus Inc. or from South Coast Biomarine and kept in flowing natural seawater tanks with aeration at 11-14 C, at the Oregon Institute of Marine Biology. Animals were fed in captivity with locally collected small mussels and blades of *Ulva*, or with minced cooked shrimp (Chuck’s Seafood). Oocyte handling, injections, and imaging were performed as described previously in detail (von Dassow et al., 2019). Briefly: oocytes were teased from chunks of ovary removed from the coelom after entry with a biopsy punch and transferred promptly to Ca2+-free artificial seawater (CaFSW); liberated oocytes were kept at 12°C and rinsed several times over the course of ~1 hr to remove follicles, then returned to microfiltered natural seawater (MFSW) at 12°C to await injection.

### *Xenopus* embryos

Adult female *Xenopus laevis* were induced to ovulate by injection with 800U HCG followed by overnight incubation at 18°C. Matured eggs were collected by gentle squeezing into 1× Modified Barth’s Saline (MBS; 88mM NaCl, 1mM KCl, 1mM MgSO_4_, 5mM HEPES, 2.5mM NaHCO_3_, 1mM CaCl_2_, pH 7.6) supplemented with high salt (5M NaCl + 0.1M CaCl_2_). Eggs were transferred to a minimal amount of 0.1× Marc’s Modified Ringers (MMR; 100mM NaCl, 2mM KCl, 2mM CaCl2, 1mM MgCl2, 5mM Hepes, pH 7.4), fertilized with macerated male *Xenopus laevis* testes, and then incubated at room temperature until fertilization envelope had risen (about 20 minutes). Fertilized embryos were dejellied in a 2% cysteine solution in 0.1× MMR and rinsed extensively before storage in 0.1× MMR. At the two-cell stage, embryos were typically microinjected with 5nl of mRNA at 0.01-1 mg ml^−1^ (needle concentration) or 5nl of protein at 8uM. For some experiments, embryos were injected a second time at the four-cell stage 2.5nl of mRNA or protein. Embryos were maintained at 18°C until imaging. For further reading on *Xenopus* cell handling, injections and imaging considerations, see (Varjabedian et al., 2018).

### *Xenopus* oocytes

Chunks of *Xenopus laevis* ovaries were removed from adult females, rinsed in 1x Barth’s solution (87.4mM NaCl, 1mM KCl, 2.4mM NaHCO3, 0.82mM MgSO4, 0.6mM NaNO3, 0.7mM CaCl2 and 10mM HEPES at pH 7.6) and then collagenase treated for 1 h at 16°C. Oocytes were rinsed extensively and allowed to recover at 16°C. Prior to imaging, stage VI oocytes were manually defolliculated and injected with 40 nl of mRNA or protein (Needle concentration: 0.01-1 mg ml^−1^ for mRNA, 1-5 μM for protein). For experiments using untagged Ect2^ΔNLS^ and RGA-3/4^WT^, mRNA encoding these proteins was either injected the night prior to imaging (Fig. 4A, Fig. S3A-D) and oocytes were incubated overnight at 16°C, or untagged Ect2^ΔNLS^ and RGA-3/4^WT^ mRNA was injected the morning of imaging and oocytes were incubated at room temperature for at least 3-5 hours prior to imaging to allow for cortical wave induction (all other figures).

### Microinjection

For starfish, microinjections were performed by transferring 150-200 oocytes, sheared recently through a narrow capillary to remove surrounding mucus, to coverslip-bottomed dishes (MatTek) pretreated with a 30-sec. rinse of 1% protamine sulfate. Oocytes were deposited in rows on a dish already emplaced on an inverted microscope with phase contrast optics, allowed to settle and bind loosely, then pressure-injected (Dagan Instruments injector and Narishige oil-hydraulic micromanipulator) with capillary glass needles (Sutter glass and P1000 puller) treated with hexamethyldisilazane. Standard injection delivers a puff slightly less than the apparent diameter of the germinal vesicle (GV), giving a nominal volume of 1-2% oocyte volume; as discussed in (von Dassow et al., 2019), the real delivery is assuredly less, but this is a visual gauge of injection volume that is reproducible from one operator to another. After injection, damaged oocytes were removed from the dish, and the remainder incubated overnight at 12-14°C.

For *Xenopus*, microinjections were performed on a PLI-100 picoinjector (Warner Instruments) with a manual micromanipulator (Narishige). Needles were pulled from capillary tubes and calibrated using oil droplets on a stage micrometer. Cells were injected in a mesh-bottomed Petri dish containing either 1× Barth’s (oocytes) or 0.1× MMR + 5% ficol (embryos). Embryos were washed after injection and put back into 0.1× MMR.

### Constructs and mRNA

All mRNA probes and constructs are contained in the pCS2+ vector. The eGFP-rGBD and mCherry-rGBD probes (to observe active Rho) as well as mCherry-UtrCH (to observe F-actin) have been described previously (Benink and Bement, 2005; Burkel et al., 2007). Plasmid DNA was linearized with a unique cutting restriction enzyme downstream of the coding sequence (usually NotI), then transcribed using the mMessage mMachine sp6 kit (Ambion). If needed, mRNA was polyadenylated using the E-PAP poly(A) tailing kit (Invitrogen). Prior to injection, RNA blends were prepared freshly from stocks with nuclease-free water (Ambion). Protein for GFP-rGBD and mCherry-UtrCH was produced and purified using a baculovirus system previously described in (Bement et al., 2015).

For starfish, the full-length RGA-3/4 clone, fused to mNeon, was a generous gift from Kuan-chung Su (Whitehead Institute, Cambridge, MA, USA). Constructs used to express wild-type purple urchin Ect2 have been described previously (Su et al., 2014); starfish Ect2 behaves similarly (not shown).

For frog, *Xenopus laevis* Ect2^ΔNLS^ was generated from Ect2^WT^ (Bement et al., 2015) by mutating the nuclear localization sequence (KRR, amino acids 379-381) to three alanines (AAA) via PCR mutagenesis. RGA-3/4^WT^ clones were generated using cDNA purchased from Horizon Discovery (catalog#: MXL1736-202797367). The cDNA was amplified via PCR and cloned into the appropriate vector backbones (empty pCS2+ or pCS2+ with C-terminal 3xeGFP) using InFusion Assembly (Takara). The GAP-dead mutant (RGA-3/4^R80E^) was generated via PCR mutagenesis. Full-length p190RhoGAP was amplified from *Xenopus* oocyte cDNA and inserted into empty pCS2+ using BamHI and XhoI restriction sites. Tagged full-length anillin (anillin-3xeGFP) was described previously in (Reyes et al., 2014). The myosin intrabody (Sf9-mNeon) was a gift from Ann Miller and described in (Hashimoto et al., 2015; Higashi et al., 2019; Nizak et al., 2003). Tagged fulllength Dia-1, Dia-2, and Dia-3 were a gift from Ann Miller and described in (Higashi et al., 2019).

### Imaging

Starfish oocytes were screened for expression level with a fluorescent dissecting scope (Leica), then selected in small groups (~20) for experiments or imaging. Oocytes were matured either in dishes by addition of 1-methyladenine (1MA) to ~10E-5 M, or by perfusion of 1MA at 10E-4 M in MFSW into imaging chambers. Perfusible imaging chambers were made crafting two ridges of vacuum grease (Dow Corning) ~1 cm apart by rolling a round toothpick (Diamond) against a Hungarian glass slide, 75×25 mm (Gold Seal), then placing no more than 20 oocytes in a drop between them, then lowering a 22×30 mm #1.5 coverslip (VWR) cross-wise to leave shelves overhanging. For some experiments these shelves are sealed by rolling a bead of vacuum grease underneath the margin, but for perfusion experiments they are left open. On an inverted microscope, perfusion is achieved by applying a paper wick (Whatman) to one shelf while adding perfusate to the other; we thus exchange 3-5 volumes of the chamber. Latrunculin B (Sigma) was dissolved freshly in MFSW from frozen aliquots at 20 mM in DMSO; vehicle concentration (0.01%) is far below the threshold (~0.5%) at which effects are detectable.

All starfish oocyte imaging was performed on an Olympus FluoView 1000 laser-scanning confocal microscope on an inverted body (Olympus IX81) using a 40x 1.15 NA superfluor water-immersion objective. This choice of objective yields a workable tradeoff between resolution, brightness, and Z-section thickness with these specimens. The confocal microscope is installed in a room kept at 16-18°C by air conditioning, hence eliminating the need for a cold stage for this work (*P. miniata* tolerates temperatures at least as high as 22°C without obvious abnormalities).

Frog oocytes and embryos were imaged using a Prairie View Laser Scanning Confocal on a Nikon Eclipse Ti base (Bruker Nano Surfaces), a Prairie View Swept Field Confocal on a Nikon Eclipse Ti base (Bruke Nano Surfaces), or an Olympus Fluoview 1000 laser-scanning confocal on an upright body (Olympus BX61WI). Data was collected using 40x 1.0 NA or 60x 1.4 NA oil objectives. All image acquisition was controlled using the Prairie View or Olympus software, respectively. Samples were mounted on custom metal slides between two #1.5 coverslips in the appropriate cell media as described in (Varjabedian et al., 2018). Cells were kept at room temperature during imaging. Early embryos were imaged starting at the 16-cell stage. Late embryos were imaged at the midblastula stage.

Light-sheet microscopy in Fig. 4F was accomplished via a custom microscope built by Jiaye He in Jan Huisken’s lab. The sample was mounted in an FEP tube sealed with an agarose plug, affixed to a rotating stage and suspended in Barth’s solution. Multidirectional illumination (Huisken et al., 2007) was done from two sides and a third objective (10×/0.3) was used for detection to capture one half of the oocyte with even illumination.

### Image Processing

All image processing was conducted using ImageJ/Fiji (Schindelin et al., 2012).

For starfish, all data are raw and unfiltered except for Fig. 2A and 2C, where background subtraction was used. Figure assembly, false coloring, and composition was conducted using Adobe Photoshop CS6, and movies were assembled and compressed using Adobe Photoshop CS6 and QuickTime Player Pro 7.

For frog, motion correction using the stack-reg plugin (Thévenaz et al., 1998) and simple image rotation was applied in Figure 3C and 3C’ to orient the furrow horizontally and correct for motion-induced artifacts. Difference subtractions (noted on figures) were performed using the Image Calculator plugin with subtraction. Kymographs were generated in Fiji by reslicing the time-lapse along a 1px-wide line drawn across the FOV at locations indicated on individual figures and using bicubic interpolation to stretch the y-axes for display. Images were pseudo-colored using custom look-up tables (LUTs) in Fiji. All figures were assembled in Adobe Illustrator, and movies were assembled using Adobe Premiere Pro.

### Image Analysis

Line profiles of signal over time were generated in Fiji using the plot Z-axis profile function on a 2-5 μm square at a representative location. The resulting measurements were imported to excel, and each signal was normalized between 0 and 1 via:

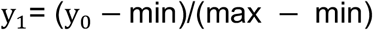

Where y_0_ represents the raw input signal, min is the minimum measured intensity, and max is the maximum measured intensity. This results in the highest peak equaling a value of 1, and the lowest minimum being equal to 0.

Wave period (auto-correlation), temporal width, relative amplitude and signal shift (cross-correlation) were initially quantified by the following process: a raw movie was divided into square boxes of size 15×15 pixels (4×4 um). Signals were spatially averaged within the boxes. This procedure resulted in the matrix J(i,j,k), i = 1,…,n; j = 1,…,m; k = 1,…,N, where N is the frame number and n,m are the numbers of boxes in vertical and horizontal dimensions. Then the autocorrelation and cross-correlation of signals were computed for each box as described previously (Landino et al., 2021). Temporal amplitudes for each box were computed as the differences between the maximal and minimal intensity values inside of the moving time window of the width approximately equal to one oscillation period as obtained from the autocorrelation curve. For each box, the average amplitude was computed by averaging over all time window positions. Temporal width was measured as the half-height width of the peaks using MATLAB function *findpeaks*. For each box, the average time width was computed by averaging over all peaks detected inside that box. Histograms, averages and standard deviations of periods, shifts between the signals, temporal amplitudes and temporal widths were computed using all boxes. For the final analysis used in the paper, a custom Python workflow described in (Swider et al., 2022) was used. Briefly, each maximum projection movie was divided up into a grid of uniform boxes. Box size was set roughly equal to wavelength, and averaged ranged from 20-25 px^2^. Wave metrics were calculated for each box, and then averaged for the entire movie. Wave period was defined as the first maxima of the auto-correlation function. Temporal width was defined as the full-width half max (FWHM) for a given wave peak. Relative amplitude was defined as the change above background, and was calculated per box as:

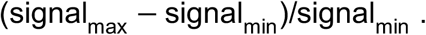

Signal shift was defined as the first maxima of the cross-correlation function.

All plots and statistical analyses were generated in Graphpad Prism 9.3.0 (www.graphpad.com). To calculate statistical significance, data was analyzed with a one-way ANOVA followed by Tukey’s post-hoc test for multiple comparisons.

The percentage of waving cells per group (Fig. S3D) was defined as the number of cells displaying any type of cortical wave activity in the field of view, divided by the total number of cells for the group, multiplied by 100. To determine end-to-end length (Fig. S3B-C), a subset of cells from each experimental group were chosen and difference subtractions were made in Fiji to enhance Rho wave segmentation. A single timepoint was chosen for each cell, and individual waves were manually segmented from beginning to end using Fiji ROIs and the segmented line tool. The lengths of the ROIs were measured in Fiji, and the average and standard deviation for each cell was calculated in Microsoft Excel.

#### Model description

We model spatiotemporal dynamics of membrane-bound active Rho (RT), inactive Rho, which consists of the membrane-bound (RD) and cytoplasmic (RDc) pools, and a fraction of the cell cortex consisting of the dynamic F-actin (F), whose polymerization is directly stimulated by active Rho. The model explicitly considers the following processes that take place on or immediately near the plasma membrane (see Fig. 7A). 1) Inactive Rho on the membrane reversibly exchanges with its cytoplasmic pool. 2) Membrane-bound inactive Rho undergoes both a low-level background activation and the Ect2-dependent positive-feedback activation. 3) Active Rho induces F-actin polymerization, e.g., via its effector Dia3 (Fig. 6G). 4) Freshly polymerized F-actin directly or via an actin-binding protein recruits Rho GAP RGA-3/4 and stimulates inactivation of Rho closing the loop of negative feedback. 5) In the wake of the diminishing Rho activity, the newly polymerized actin disassembles and recycles back to the cytoplasm. These biochemical reactions and the diffusion of species are described by the following system of reaction-diffusion equations:

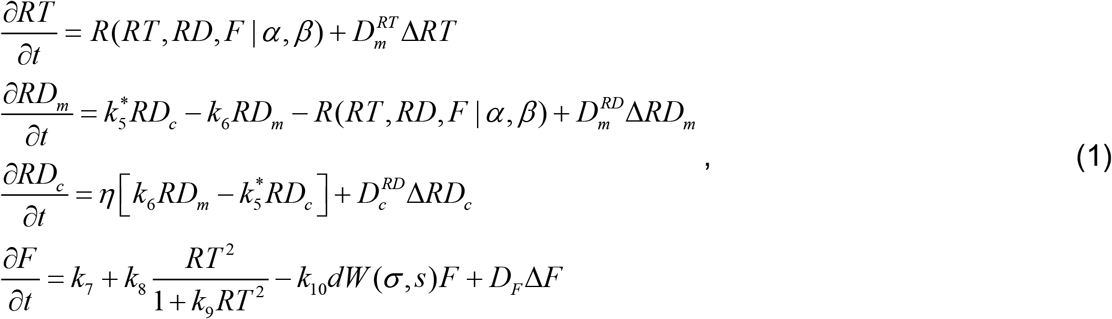

where *R* is the reaction function:

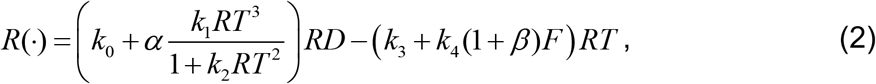

and *α*, *β* are the non-dimensional effective protein concentrations of Ect2 and RGA-3/4, respectively. In the order of presentation, the terms of the reaction function describe the rates of the low-level background activation of Rho, autocatalytic Ect2-dependent positive-feedback activation of Rho, constitutive inactivation of Rho, and, finally, the F-actin-dependent inactivation of Rho. In (2) we assume that F-actin exhibits some inhibitory effect on the activity of Rho even when the concentration of RGA-3/4 is zero.

This provides for the possibility that other actin-binding proteins (or the polymer itself) contribute to the reported inhibitory effect of F-actin on Rho activity (Bement et al., 2015). This assumption increases the flexibility of model (1) without changing its qualitative behavior. Importantly, the model values of Ect2 and RGA-3/4 concentrations are determined up to the unknown scaling factors, assuming, in the simplest scenario, a linear dependence between the injected amounts and the resulting cellular concentrations of proteins. A possibility of the nonlinear dependence, however, cannot be ruled out, especially in the case of RGA-3/4, which is injected as mRNA. Thus, the efficiency of mRNA translation could be potentially a saturable function of the amount of injected mRNA.

A stochastic noise term *dW*(*σ,s*) in (1) represents multiple unaccounted biochemical processes that contribute to the local rate of F-actin depolymerization. This choice is strongly motivated by the inherent complexity of regulation of F-actin dynamics, which is modulated by a large number of diverse actin-binding proteins (Pollard, 2016). In our model, *dW*(*σ,s*) is a spatially correlated Gaussian stochastic field with spatial correlation *s*, standard deviation *σ* and the mean value 1. The stochastic field was randomly generated every *f* seconds throughout the entire model simulation.

Model (1) does not account for Rho production or degradation and, therefore, conserves the total cellular amount of the GTPase. The exchange between the membrane-bound and the cytoplasmic pools is governed by a small parameter *η*, which represents the ratio between the volumes of the notional membrane and cytoplasmic compartments and is inversely proportional to the cell radius. Assuming for clarity that all biochemical processes modeled by us as happening on the membrane take place in a 100 *nm* thick layer of the cytoplasm above the surface of lipid membrane, we estimate *η_f_* ≈ 5 ·10^−5^ for the frog oocyte with the characteristic diameter of 1.2 *mm*. For comparison, in the budding yeast cell, in which mass-conservation of another small GTPase Cdc42 is essential to ensure the uniqueness of the bud (Goryachev and Pokhilko, 2008), this ratio is nearly three orders of magnitude larger, *η_y_* ≈ 0.03. Thus, for the large cells of oocytes and embryos, it is justifiable to assume that the entire membrane-bound amount is only a small fraction of the total cellular quantity of the GTPase and take a limit *η* → 0 in (1).

Under this assumption, *RD_c_* = *RD*_0_ and the model (1) reduces to

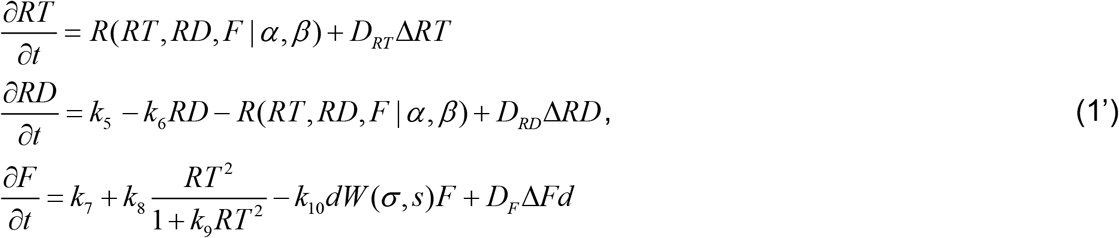

where, 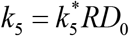, and 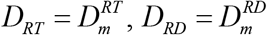. Since both diffusion coefficients describe the membrane-bound active and inactive forms of the GTPase, they no longer differ by orders of magnitude as in the models postulating that RT and RD are present on the membrane and in the cytoplasm, respectively (Holmes et al., 2012). Instead, we assume that membrane diffusion of active Rho is only mildly slower than that of the inactive GTPase, *D_RT_* < *D_RD_*, which is believed to be caused by the interaction of RT with its multiple effectors (Hodge and Ridley, 2016). We also increase the flexibility of our model by allowing *D_F_* > 0. The non-vanishing diffusivity of the inhibitor in our model could be interpreted as both the reversible binding of a Rho-inhibitory molecule, such as RGA-3/4, to static F-actin, and as the dynamic rearrangement of F-actin due to the processes of polymer breaking and re-annealing (Pollard and Craig, 1982). To facilitate the comparison of simulated and experimental waves, we scaled the model parameters to match the time period and the wavelength of the waves observed in the immature frog oocytes.

Model (1’) was numerically simulated with finite difference method on 2D spatial domains with periodic boundary conditions using a custom-built C code. The simulations were initiated with the concentrations of the parameter-specific uniform steady state. Linear stability analysis of the model steady state was performed numerically in 1D using Mathematica.

#### Calculation of the histogram for Figure 8F

84 experimental movies belonging to the six groups of experiments, grouped by the concentration of injected RGA-3/4 mRNA (0 ng/μL, n = 10; 33 ng/μL, n = 10; 66 ng/μL, n = 13; 166 ng/μL, n = 24; 333 ng/μL, n = 17; 1000 ng/μL, n = 10) were scored by the presence (1) or absence (0) of morphological features from the ten categories defined below. To compute the histogram for each concentration group, the sum of occurrences of each feature calculated for each concentration of injected RGA-3/4 mRNA was divided by the total sum of occurrences across all experiments. Ten characteristic morphological features were defined as follows:

1. Rho flickers – spatially disorganized local flashes of Rho activity with no evidence of spatial propagation.
2. Turbulence 1 – individual maxima of Rho activity randomly emerge and erratically move a short distance prior to disappearance. Propagation is evident from the kymograph.
3. Turbulence 2 – fragments of waves are well formed and densely populate the field of view occasionally forming short-lived, localized wave trains. Spatial correlation is absent, while temporal correlation is clearly detectable in the kymograph.
4. Turbulence 3 – turbulent dynamics similar to T2 but detectable within spatially limited areas, which are surrounded by the well-organized wave trains and spirals.
5. Lamellar wave trains – persistent domains of flat or slightly curved waves with robust propagation.
6. Dislocations of wave fronts.
7. Grain boundaries – pattern emerging at the interface of two wave trains with near orthogonal wave vectors.
8. Single armed spiral waves – well-formed spiral waves containing at least 2-3 full turns.
9. Two-armed spiral waves – stable two-armed spiral waves containing at least 2-3 full turns.
10. Line defects – boundaries between the wave trains with the angle between their wave vectors close to 180 degrees (antiparallel wave vectors).

## Reproducibility of experiments

When possible, all results reported come from at least three independent experiments using biologically distinct cells to eliminate batch-effects. The specific number of cells and experiments are indicated in each figure legend.

## Summary of supplemental material

Supplemental Figure 1 contains additional starfish data, Supplemental Figures 2–4 contain additional frog data, Supplemental Videos 1-4 correspond to starfish figures, and Supplemental Videos 5-11 correspond to frog figures.

## Acknowledgements

We thank Ann Miller for sharing the Sf9-mNeon and Dia1, Dia2 and Dia3 clones, Kuan-chung Su for the starfish RGA-3/4 clone, and Andie Bolton for her help with general cloning and reagent preparation. We would also like to thank Kevin Sonnemann for preparing and purifying the GFP-rGBD and mCherry-UtrCH protein used in this study. This work was supported by NSF grant MCB-1614190 (W.M.B.) and MCB-1614606 (G.v.D) the NIH grant RO1GM052932 (W.M.B) and the BBSRC grants BB/P006507 and BB/P01190X (A.B.G.). The authors declare no competing financial interests.

## Author contributions

**Ani Michaud:** Conceptualization, data curation, formal analysis, investigation, methodology, software, validation, visualization, writing (original draft), writing (review and editing)

**Marcin Leda:** Conceptualization, data curation, formal analysis, methodology, visualization, writing (review and editing)

**Zachary T. Swider:** Investigation, methodology, software, validation, writing (review and editing)

**Songeun Kim:** Investigation, methodology, validation, writing (review and editing)

**Jiaye He:** Investigation, methodology, resources, writing (review and editing)

**Jennifer Landino:** Conceptualization, resources, writing (review and editing)

**Jenna R. Valley:** Investigation, data curation, resources, validation

**Jan Huisken:** Methodology, resources, writing (review and editing)

**Andrew B. Goryachev:** Conceptualization, data curation, formal analysis, funding acquisition, methodology, software, resources, supervision, validation, visualization, writing (original draft), writing (review and editing)

**George von Dassow:** Conceptualization, data curation, formal analysis, funding acquisition, investigation, methodology, resources, supervision, validation, visualization, writing (original draft), writing (review and editing)

**William Bement:** Conceptualization, data curation, funding acquisition, investigation, methodology, resources, supervision, validation, visualization, writing (original draft), writing (review and editing)

